# Morphospace analysis reveals divergent cellular behaviours driving tissue internalisation during insect gastrulation

**DOI:** 10.64898/2026.06.30.735526

**Authors:** Margherita Battistara, Wolfram Pönisch, Matthew A. Benton, Ewa K. Paluch

**Affiliations:** Department of Physiology, Development and Neuroscience, University of Cambridge, Cambridge CB2 3DY, UK; Department of Zoology, University of Cambridge, Cambridge CB2 3EJ, UK

## Abstract

During gastrulation, embryonic epithelia undergo large-scale remodelling to internalise the mesoderm. Whether the cellular mechanisms underlying these tissue deformations are conserved across species remains unclear. Using quantitative cell-shape embedding, we compare cellular behaviours underlying gastrulation in two insects. We show that while *Drosophila* mesoderm cells follow a coordinated shape change sequence during gastrulation, mesoderm cells in the beetle *Tribolium* display heterogeneous shapes with no clear trajectory. This heterogeneity arises from two distinct internalisation modes: canonical apical constriction-driven tissue invagination, and early internalisation through out-of-plane cell divisions. Interpretable machine learning together with *in vivo* perturbations identify nuclear crowding as a key predictor of out-of-plane divisions and show that, in both species, proliferation slows tissue folding while promoting individual cell internalisation. Our study showcases morphospace analysis as a powerful tool for dissecting the cellular basis of tissue morphogenesis and reveals a conserved antagonistic relationship between cell divisions and tissue folding during gastrulation.

## INTRODUCTION

Embryonic development proceeds via a series of precisely ordered and coordinated cellular behaviours^1,2^. Understanding how individual cell behaviours drive global tissue morphogenesis, and how those behaviours have evolved to give rise to the diversity of life on earth, remains a central challenge in biology.

Gastrulation is the first large-scale coordinated tissue rearrangement during animal embryogenesis, during which the mesodermal and endodermal precursors internalise from a simple, generally monolayered blastoderm epithelium^3^. Despite the deep evolutionary origin of gastrulation, the cellular mechanisms driving tissue internalisation exhibit extreme diversity across species^3,4^. Indeed, while gastrulation generally involves a combination of apical constrictions and loss of epithelial polarity / dismantling of adherens junctions associated with epithelial-to-mesenchymal transition (EMT) of mesoderm and endoderm cells^5^, the timing and coordination of these cellular behaviours greatly vary between species^6^. The resulting modes of tissue internalisation span a continuum from globally coordinated tissue bending or folding, to internalisation through individual cell ingressions.

For instance, in chick and mouse embryos, collective apical constriction first generates a shallow invagination, the primitive streak, where mesoderm cells then gradually disassemble their apical connections and ingress individually^7–9^. Individual cell ingression also drives mesoderm and/or endoderm internalisation in numerous invertebrate species, including examples from cnidarians^10^ and sea urchins^11^. In contrast, other cnidarians^12^ and insects such as the fruit-fly *Drosophila melanogaster* (hereafter *Drosophila*) internalise tissue via coordinated epithelial sheet bending and invagination. Finally, many animal embryos, especially reptiles, fish and amphibians, combine features of cell ingression and epithelial invagination with tissue involution in mixed internalisation modes^16–19^.

The cellular mechanisms driving mesoderm internalisation have been particularly well described in *Drosophila*^20,21^, making it the canonical example of epithelia invagination^22^. Starting from a monolayered epithelium, columnar blastoderm cells synchronously elongate and apically constrict, causing the formation of a ventral furrow. This leads to mesoderm tissue folding inwards as a coherent monolayer. Gastrulation in other insects is highly diverse, ranging from single cell ingression^23^ to internalisation of a rigid epithelial sheet without bending^24^, yet detailed analysis of the underlying cellular behaviours is mostly lacking. The flour beetle *Tribolium castaneum* is the best studied insect model beside *Drosophila*^25–27^. *Tribolium* mesoderm gastrulation involves homologues of the same key regulators as *Drosophila*, including *Tc-twist* (*Tc-twi*)*, Tc-snail* (*Tc-sna*), and the Fog signalling pathway, and is also driven by tissue bending and invagination of a ventral furrow^28–31^. However, this folding is more variable across the embryo^29^, and cellular behaviours other than apical constriction, including mesoderm cell divisions and individual cell ingressions through EMT, have been suggested to play a role in facilitating mesoderm internalisation^29^. The relative importance of such individual behaviours versus coordinated furrow invagination in *Tribolium* mesoderm internalisation remains unclear.

Here, we combine quantitative cell shape mapping with machine-learning analysis of tissue dynamics and perturbation experiments, to dissect the cellular behaviours underlying mesoderm gastrulation in *Drosophila* and *Tribolium*. Our findings uncover an unexpected mechanistic diversity underlying conserved gastrulation outcomes and suggest that evolutionary modulation of the balance between proliferation and apical constriction may have shaped distinct modes of insect mesoderm invagination.

## RESULTS

### Morphospace analysis reveals distinct cell shape trajectories during mesoderm gastrulation in *Drosophila* and *Tribolium*

We used morphospace analysis^32^ to quantitatively characterise the cellular shape changes underlying mesoderm gastrulation in *Drosophila* and *Tribolium* (Fig. 1). We collected and fixed *Drosophila* and *Tribolium* embryos at regular time-points during gastrulation. We then performed fluorescent staining of nuclei with DAPI and cell outlines with phalloidin (Fig. 1A), combined with hybridisation chain reaction (HCR) staining to detect expression of the mesoderm patterning genes *twi* and *sna* (*Tc-twi* and *Tc-sna* in *Tribolium*), identifying presumptive mesodermal cells (Fig. S1A). *Tribolium* gastrulation has been reported to vary along the anterior-posterior axis^29^, and we therefore used the expression of the segmentation gene *Tc-even-skipped*^33^ to compare similar anterior regions in different embryos. We then segmented mesoderm cells in 3D in the *Drosophila* and *Tribolium* embryos (Fig. 1A, see Methods). We analysed cell shapes in 12 *Drosophila* fixed embryos staged at 6 time-points (Fig. 1C, C’) and 14 *Tribolium* fixed embryos at 7 time-points (Fig. 1D, D’) during mesoderm gastrulation, obtaining two embryos per time-point from the blastoderm stage up until mesodermal spreading. Our analysis yielded a total of 3,794 *Drosophila* and 4,075 *Tribolium* 3D cell contours.

**Figure 1.**
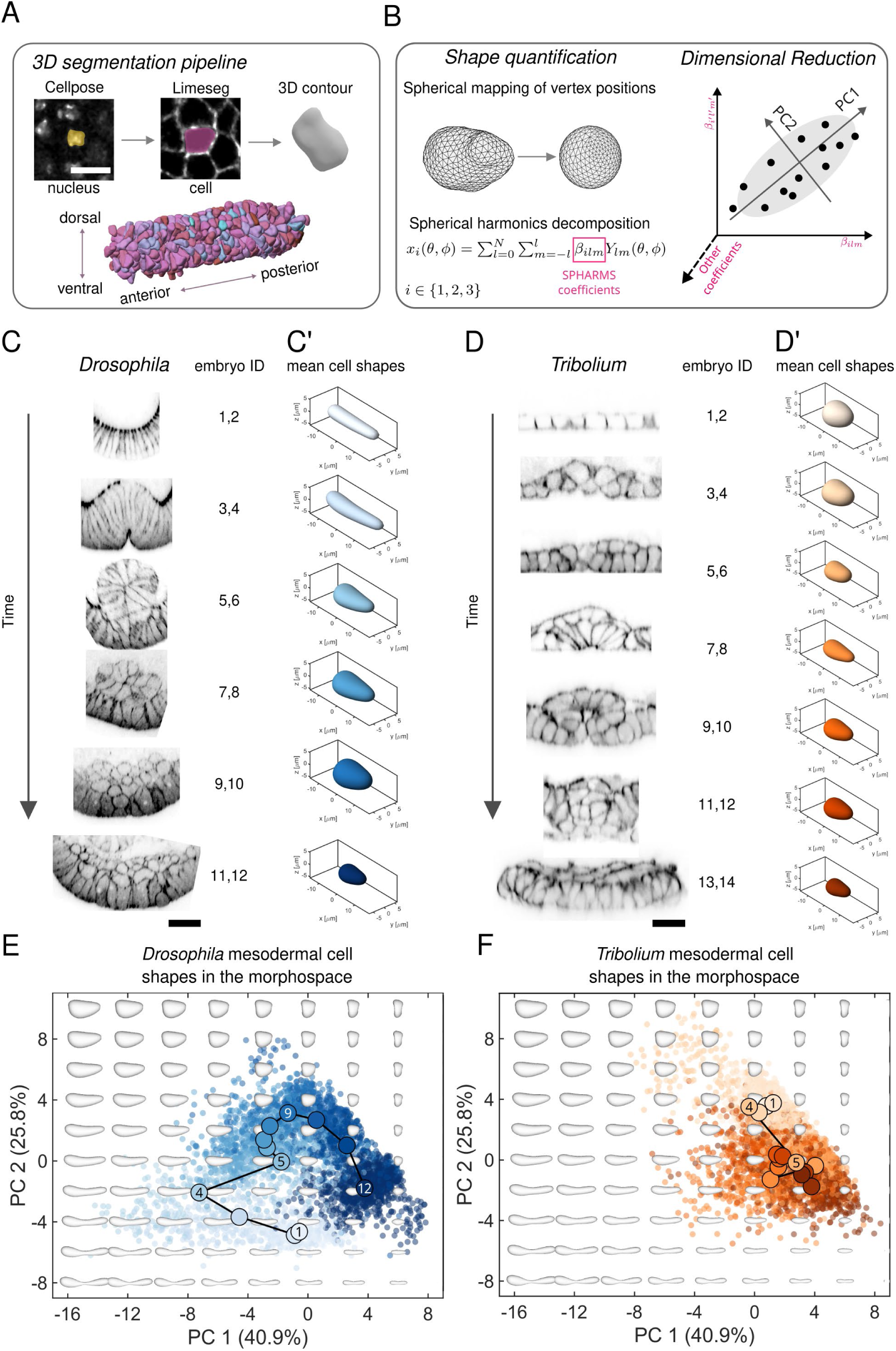
Morphospace analysis of cell shape trajectories during mesoderm gastrulation in *Drosophila* and *Tribolium*. (A) Mesodermal cell segmentation in 3D. Top: segmentation workflow. Confocal image of nuclei (DAPI) with overlaid pink masks representing segmented nuclear contours obtained using Cellpose. Nuclear centroid coordinates were extracted and used as seeds for 3D cell segmentation based on the phalloidin-labelled cell outline using LimeSeg. For each nucleus, a spherical seed was initialised and iteratively expanded to reconstruct the full 3D cell contour. Bottom: whole-tissue 3D rendering of segmented mesodermal cells. Scale bar: 10 µm. (B) Outline of the morphospace analysis pipeline. Segmented cell contours are decomposed into spherical harmonics (SPHARMS) coefficients (left). The large SPHARMS coefficients dataset is dimensionality reduced using PCA to generate a cell shape morphospace (right). (C and D) Cross-sections of the ventral side of *Drosophila* (C) and *Tribolium* (D) embryos stained with phalloidin during mesoderm development; embryos were arranged in time based on overall tissue morphology. Scale bar: 20 µm. (C’ and D’) Mean 3D mesodermal cell contours corresponding to the developmental stages of the embryos in (C) and (D). The mean shapes were reconstructed by averaging the spherical harmonics coefficients of all segmented mesoderm cells from two embryos per developmental stage. (E and F) Mesoderm 3D cell shapes from *Drosophila* (E) and *Tribolium* (F) embryos undergoing gastrulation, plotted in the PC1-PC2 morphospace. The morphospace was built using all segmented mesoderm cell contours from all analysed *Tribolium* and *Drosophila* embryos. Background greyscale shapes: reconstructed theoretical cell shapes for different combinations of PC1 and PC2 in the morphospace. Data-points: individual cell shapes: in (E), shades of blue correspond to cells from different Drosophila embryos; in (F), shades of orange correspond to cells from different Tribolium embryos. Darker tones represent later time points, as in (C’) and (D’). Large dots: centres of mass of the distributions for each individual embryo; numbers correspond to embryo numbers in (C) and (D). The black lines highlight the mean mesoderm cell shape trajectory in the morphospace.

We then adapted a morphospace analysis pipeline we recently developed^32^ to quantitatively analyse our cell contour dataset (Fig. 1B). We first parametrised each 3D cell contour using spherical harmonics (SPHARMS) decomposition, yielding a list of coefficients that quantitatively describe the cell contours. We found that 40 SPHARMs, corresponding to 5043 SPHARM coefficients per cell were sufficient to describe the cell shapes in our dataset with a high degree of accuracy (see Methods and Fig. S1B). We then used principal component analysis (PCA) dimensionality reduction to analyse this SPHARMs coefficients dataset. We found that the first two principal components (PCs) of shape variation together account for 66.7% of shape variability across our cell contour dataset, while each subsequent PC represents less than 5% of shape variability (Fig. S1C). We thus focused on PC1 and PC2 of shape variation for further analysis. Increasing PC1 represents a transition from elongated, large and wedge-shaped to compact, smaller morphologies, while increasing PC2 captures a shift from elongated to rounder shapes (Fig. S1D). We then used PC1 and PC2 as morphospace axes to analyse the cell shape changes associated with mesoderm invagination in *Drosophila* and *Tribolium* gastrulation.

Morphospace analysis of *Drosophila* cell shapes revealed a clear ensemble shape trajectory (Figs. 1E and S1E, solid lines highlight shape trajectory). Initially, cells displayed columnar morphologies, which then elongated, increased in volume and became increasingly wedge-shaped, reflected by a steep decrease in PC1 (Fig. 1E, compare embryos 1 and 2 with embryos 3 and 4). After tissue invagination (Fig. 1E, embryo 5 onwards), cells adopted shorter and rounder morphologies, corresponding to an increase in both PC1 and PC2. Finally, mesodermal cells displayed progressively smaller and spread morphologies, reflected by a decrease in PC2 and a further increase in PC1 (Fig. 1E, embryos 9-12). Together, our morphospace analysis provides a quantitative description of the 3D shape changes displayed by mesoderm cells during *Drosophila* gastrulation.

In contrast to *Drosophila*, morphospace analysis of *Tribolium* mesoderm cells did not reveal a clear shape trajectory during gastrulation (Figs. 1F and S1F). Instead, cells exhibited a sharp transition between two general shapes. At early stages, *Tribolium* mesodermal cells displayed mostly cuboid morphologies (Fig. 1F, embryos 1-4), corresponding to higher values of PC2 compared to early-stage *Drosophila* mesoderm columnar cells (Fig. 1F, embryos 1 and 2). Between embryos 4 and 5, *Tribolium* mesoderm cells transitioned towards smaller and overall more elongated shapes, reflected by a decrease in PC2 (Fig. 1F). Differences in cell shapes between successive stages of tissue internalisation however could not be well separated when considering mean cellular shapes (Fig. 1F, embryo 5 onwards). To further explore the dataset, we applied an alternative non-linear dimensionality reduction algorithm, t-distributed Stochastic Neighbour Embedding (t-SNE), which embeds high-dimensional data into two dimensions by rearranging data-points according to local similarities in morphology, and thus helps visually group cells with similar shapes. However, we found that, like PCA, t-SNE analysis did not yield a clear mesoderm cell shape trajectory during *Tribolium* gastrulation (Fig. S1G, H).

Together, our cell contour dataset and morphospace analysis constitute a rich quantitative resource characterising 3D cell shape changes during insect gastrulation. This quantitative analysis shows that despite similar patterns of mesodermal gene expression (Fig. S1A) and similar overall tissue shape changes (Fig. 1C, D), mesoderm cell shape dynamics significantly differ between *Drosophila* and *Tribolium* gastrulation. While in *Drosophila* mesoderm cells display consistent shapes at each stage of gastrulation resulting in a well-defined mean shape trajectory, in *Tribolium* shapes do not display a clear trajectory.

### *Tribolium* gastrulation is associated with early internalisation of a subset of cells

We then further investigated the cellular shape dynamics underlying *Tribolium* mesoderm gastrulation. We first asked to what extent cell shape variability within individual embryos might mask directed shape changes in the *Tribolium* mesoderm morphospace. To assess shape variability, we first confirmed that the distance between points in the morphospace quantitatively reflects differences between actual cellular shapes (Fig. S2A). Using morphospace distance (across all PCs, see Methods) to quantify cell shape differences, we then computed the total variance of mesoderm cell shapes for each embryo (Fig. S2B). We found that embryos 3 and 4 displayed the highest cell shape variance in our dataset. In contrast to *Drosophila*, where cell division is suppressed from cellularisation until after gastrulation^34–36^, *Tribolium* mesoderm cells undergo a wave of divisions during gastrulation^37^. As cell divisions in early embryos are typically associated with an abrupt drop in cellular volume between the mother and daughter cells, we wondered whether division-associated volume variations might account for enhanced cell shape variability during *Tribolium* gastrulation. We thus reanalysed *Tribolium* mesoderm cell shape trajectories after normalising each cell for volume by building a new morphospace. This analysis did not reveal any clearer cell shape trajectory (Figs. 2A and S2C). However, cell shape variance of volume-normalised shapes now peaked at the time of ventral furrow formation (Fig. 2B embryos 7 and 8, and Fig. S2C), indicating that mesoderm cells display enhanced variability in cellular shapes, independent of division-associated volume changes, at this stage of gastrulation.

**Figure 2.**
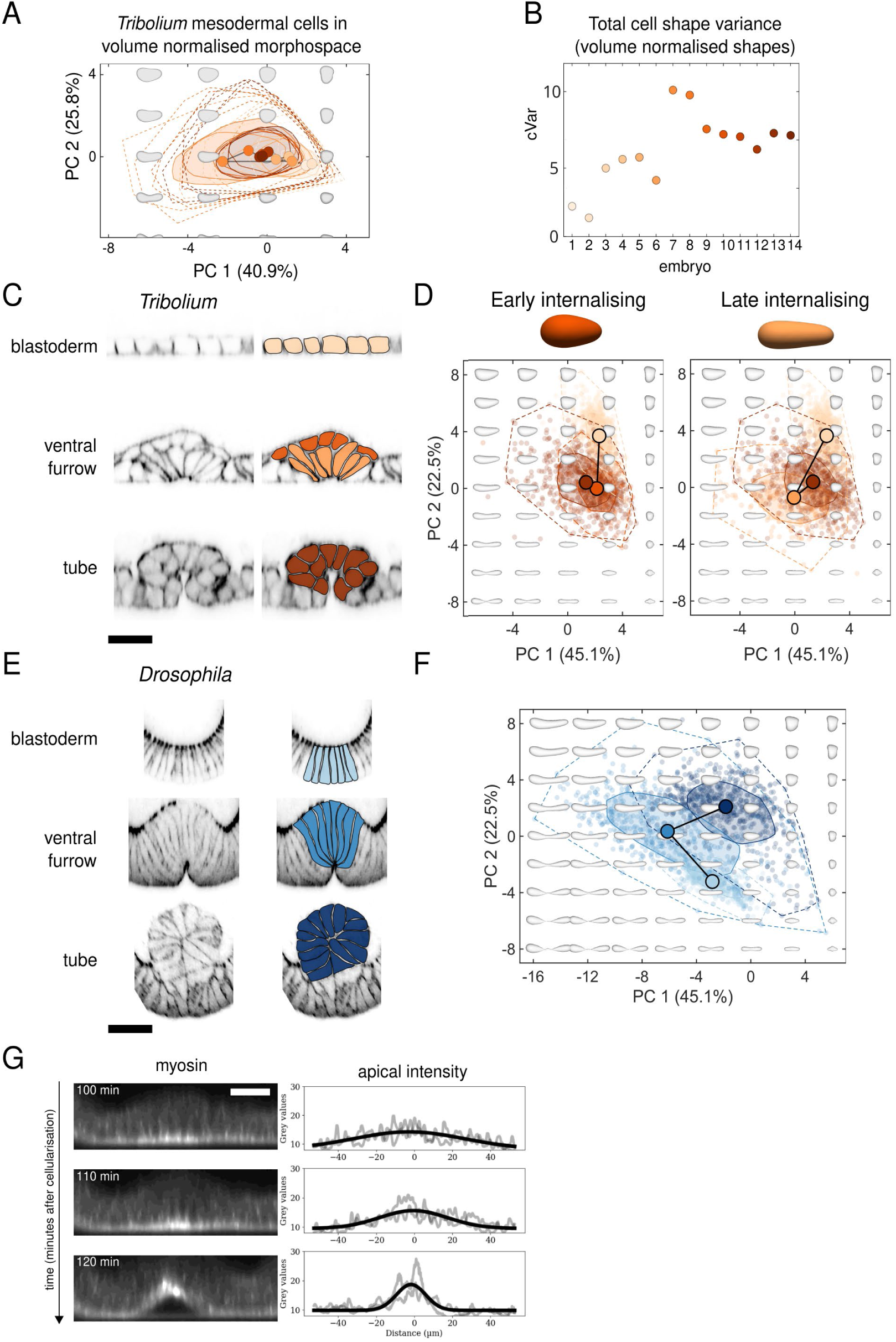
*Tribolium* gastrulation is associated with early internalisation of a subset of cells. (A) Bag-plot representation of mesoderm 3D cell shapes from *Tribolium* embryos undergoing gastrulation, plotted in the PC1-PC2 morphospace built after volume normalisation. Polygons represent bagplots: the inner polygon (bag) contains 50% of points, the outer polygon (fence) corresponds to the convex hull of all data. Black line: cell shape trajectory as defined in Fig. 1E,F. (B) Total variance of all spherical harmonics functions for each stage after cell volume normalisation (see Eq. 2 and Methods). (C) Cross-sections of the ventral side of *Tribolium* embryos stained with phalloidin during three successive stages of mesoderm invagination (blastoderm, ventral furrow and tube stages), segmented mesoderm cells are colour-shaded on the right, highlighting two cell populations at ventral furrow stage: cells displaying apical constriction (light orange) and cells displaying early internalisation (dark orange). Scale bar, 20 µm. (D) Mesoderm cell shapes from embryos at stages depicted in (C) plotted in the PC1-PC2 focused morphospace, showing distinct cell shape trajectories for cells internalising early and for apically constricting cells. The morphospace was built using all segmented mesoderm cell contours from *Tribolium* and *Drosophila* embryos at the blastoderm, ventral furrow and tube developmental stages (2 embryos per stage). Background greyscale shapes: reconstructed theoretical cell shapes for different combinations of PC1 and PC2. Data-points: individual cell shapes; the shades of orange correspond to developmental stages highlighted in (C). Polygons represent bagplots: the inner polygon (bag) contains ≤50% of points, the outer polygon (fence) corresponds to the convex hull of all data. Large dots: centre of mass of the distribution for each individual embryo. The black lines highlight the mean morphospace cell shape trajectories. (E) Cross-sections of the ventral side of *Drosophila* embryos stained with phalloidin during three successive steps of mesoderm invagination (blastoderm, ventral furrow and tube stages), segmented mesoderm cells are highlighted in blue (right). Scale bar, 20 µm. (F) Mesoderm cell shapes from embryos at stages depicted in (E) plotted in the same PC1-PC2 focused morphospace as displayed in (D). Data-points and trajectory meaning: as in (D). (G) Left: Time-lapse of the ventral side of a gastrulating *Tribolium* embryo transiently expressing squash-GFP to mark myosin. Scale bar, 20 µm. Right: quantification of apical myosin intensity. The myosin profile along the ventral cross-section in two embryos is displayed (greyscale lines) as well as a Gaussian fit to their mean (black line).

Interestingly, we also observed that at ventral furrow formation stage in *Tribolium*, a subset of mesodermal cells appears already internalised while others are wedge-shaped (Fig. 2C), in agreement with past work that showed that the *Tribolium* mesoderm appears multi-layered during gastrulation^29^. We thus asked whether these two cell populations occupied distinct locations in the morphospace, which could account for overall enhanced cell shape variability at this stage. To this end, we constructed a minimal morphospace focusing on embryos fixed before, during and after ventral furrow formation (Figs. 2C and S3A,B). These stages correspond to the blastoderm stage, the ventral furrow formation *per se* and the tube stage. When we separated out the mesodermal cells undergoing early internalisation (dark orange in Fig. 2C and 2D) from the other mesodermal cells (light orange in Fig. 2C and 2D), we found that the two cell populations displayed distinct morphospace trajectories (Fig. 2D and S3C). Specifically, early internalising cells displayed shortening and rounding, characterised by a strong decrease in PC2, at the ventral furrow formation stage and did not significantly change morphology at the tube stage (Fig. 2D left). In contrast, late internalising cells displayed a combined decrease in PC1 and PC2 at ventral furrow formation, reflecting cell elongation and apical constriction, prior to shortening and rounding at tube formation, reflected by a slight increase in both PC1 and PC2 (Fig. 2D right). The shape trajectory of the late internalising cells resembled the shape behaviour of *Drosophila* mesoderm cells at similar stages of tissue invagination (Figs. 2E,F and S3D), albeit with a different starting point since *Tribolium* blastoderm cells are cuboidal rather than columnar like in *Drosophila* (compare blastoderm stage cells in Fig. 2C and 2E). In *Drosophila*, apical constrictions driving ventral furrow formation are powered by myosin-driven contractions^38,39^. We thus analysed myosin distribution in the *Tribolium* mesoderm via live imaging of embryos transiently expressing the non-muscle myosin II regulatory light chain (hereafter referred to as myosin) fused with GFP (squash-GFP), and found that myosin accumulated apically prior to ventral furrow formation (Fig. 2G, Video S1). This strongly suggests that apically constricted cell shapes in the late internalising *Tribolium* mesoderm result from myosin-driven apical contractions, as described in *Drosophila*.

Taken together, our quantitative shape analysis identifies ventral furrow formation as the time-point of highest cell shape variability, aside from division-induced volume changes, during *Tribolium* gastrulation. Focused analysis of cell shape dynamics shows that two distinct morphospace cell trajectories co-exist during tissue invagination in *Tribolium*: while late internalising cells display myosin-driven apical constriction similarly to their *Drosophila* counterparts, a sub-population of mesoderm cells internalise earlier and do not display further shape changes following internalisation.

### Out-of-plane divisions and cell extrusion drive early cell internalisation in the *Tribolium* mesoderm

We then asked what cellular behaviours drive the early cell internalisations observed during *Tribolium* gastrulation. We first hypothesised that early internalisation might be driven by basal cell extrusion, a common mechanism of epithelial cell delamination^40–42^. This is in line with previous suggestions that early internalisation might result from early EMT of posterior mesoderm cells^29^. We thus assessed individual cell dynamics using live imaging of gastrulating *Tribolium* embryos transiently expressing fluorescently labelled histone H2A (to mark DNA) and a membrane label (Gap43-mNeongreen) (Fig. 3A and Video S2). While we did observe basal cell extrusion (Fig. 3B), it was displayed by only 5% of mesoderm cells and was thus unlikely to account for the relatively high numbers of early internalised cells we observed (Fig. 2D). However, we noticed that in mesoderm cells undergoing divisions the division axis was not always oriented within the plane of the epithelium as is typical during cell division in epithelial monolayers^43^, but was sometimes oriented perpendicular to the epithelial plane (Figs. 3C and S4A). Our live analysis showed that when mesodermal cells divided perpendicular to the epithelial plane, the basally positioned daughter cell left the epithelium and internalised early (Video S1). Overall, we found that over the course of *Tribolium* gastrulation, 25% of mesodermal cell divisions occurred out-of-plane and resulted in early internalisation of one of the daughter cells (Fig. 3C). Together, these findings show that early internalisation of *Tribolium* mesodermal cells is primarily driven by out-of-plane divisions, with a small contribution from cell extrusion.

**Figure 3.**
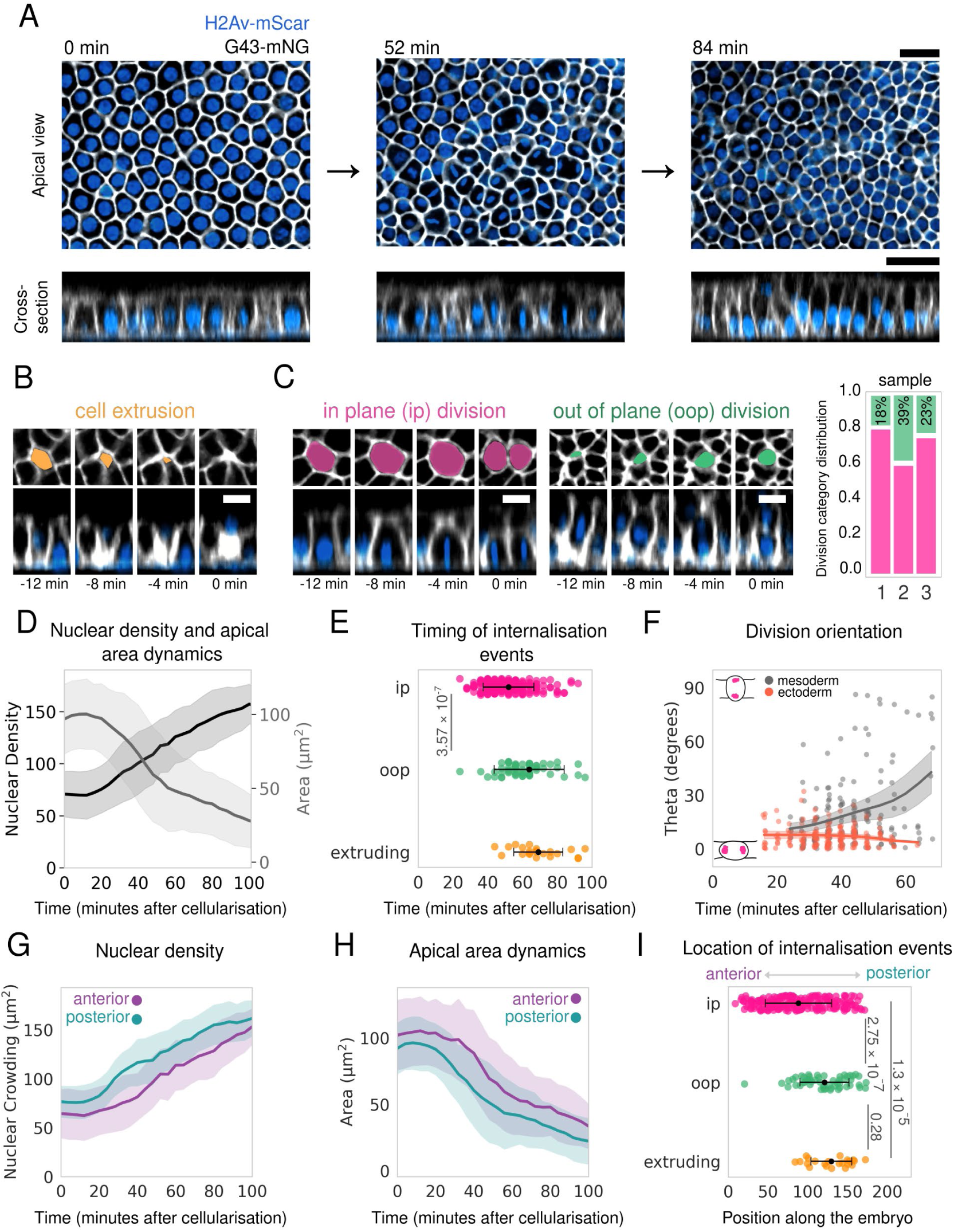
Individual cell internalisation in the *Tribolium* mesoderm is mediated by out-of-plane divisions and cell extrusions. (A) Confocal microscopy time-lapse of the *Tribolium* mesoderm in an embryo transiently expressing H2Av-mScarlet to label nuclei (blue) and G43-mNG to label the plasma membrane (greyscale). Top: apical view (average projection of grazing section of ∼4µm). Bottom: cross-section through the middle of the top panels, obtained by averaging over ∼4µm. Scale bar, 20 µm. (B) Representative time-lapse images of the *Tribolium* mesoderm showing a cell extruding in both apical (top) and cross-sectional (side) orientations. The apical view highlights change in the cell’s surface area, while the cross-sectional view shows the dynamic positioning of cells over time at key time points. Time zero corresponds to the time in which the cell extrudes. Scale bar, 10 µm. (C) Left: representative time-lapse images of the *Tribolium* mesoderm in both apical (top) and cross-sectional (side) orientations. Cells are seen undergoing in-plane division (magenta) and out-of-plane division (green). Time zero corresponds to the time in which the cell divides. Scale bar, 10 µm. To the right, stacked plot quantifying division category in three embryos. The percentage of out of plane divisions out of the total of division events are reported. (D) Nuclear density (black) and cell apical area (greyscale) over time. Lines represent mean values calculated from cells tracked in *N* = 3 embryos (*n =* 170 mother and *n* = 235 daughter cells tracked); the shaded area represents the standard deviation. (E) Scatter plot of the timing of in plane division (pink, *n* = 189 events) and out-of-plane divisions (green, *n* = 61), and the timing of cell extrusions (orange, *n* = 22), occurring within 100 minutes after cellularisation. *N* = 3 embryos were analysed. The Mann-Whitney p-value is reported. (F) Scatter plot of division angles in the mesoderm and ectoderm with respect to the embryo surface as a function of time after cellularisation. Data-points: angle (theta) between the apical surface and the line connecting the two daughter cell nuclei in the first frame after cytokinesis. *n* = 162 divisions in the ectoderm; 169 divisions in the mesoderm, in *N* = 3 embryos. Solid lines: mean division orientation, shaded area: standard error of the mean. (G,H) Nuclear density (G) and cell area (H) as a function of time after cellularisation, in the anterior (purple, *n* = 119 mother cells, 171 daughter cells) and posterior (teal, *n* = 131 mother cells, *n* = 197 cells) regions of the embryo. *N* = 3 embryos were analysed. Lines: mean values; shaded areas: standard deviation. (I) Scatter plot of the localisation of in-plane divisions (pink, *n* = 189 events), out-of-plane divisions (green, *n* = 61), and cell extrusions (orange, *n* = 22) along the anteroposterior axis, given in µm. *N* = 3 embryos were analysed. The Mann-Whitney p-value is reported.

### Early cell internalisation events during *Tribolium* gastrulation predominantly occur in regions of highest tissue crowding

We then investigated what causes the out-of-plane divisions in the *Tribolium* mesoderm. Past work in other systems points to epithelial tissue crowding as a major driver of both cell extrusion and division out of the epithelial plane^44–46^. The *Tribolium* mesoderm is apically constricting and proliferating, which could progressively increase tissue crowding. Indeed, we found that nuclear density increased and cell apical area decreased over time in the gastrulating mesoderm, indicating increasing tissue crowding (Fig. 3D). Both out-of-plane divisions and cell extrusions occurred on average later than in-plane divisions (Fig. 3E), correlating with increased tissue crowding. Furthermore, spindle orientation in the mesoderm became increasingly variable and deviated from in plane orientation through gastrulation (Fig. 3F). Collectively, these findings show that increasing mesoderm tissue crowding correlates with progressive increase in out-of-plane divisions and cell extrusions.

The posterior of the *Tribolium* embryo folds earlier and more deeply than the anterior^29^ (Fig. S4B and C), prompting us to compare tissue crowding and cell internalisation frequency between these two regions. We found that mean apical cell area was smaller and nuclear density was larger in the posterior versus anterior half of the mesoderm throughout gastrulation (Fig. 3G,H). Interestingly, both out-of-plane divisions and cell extrusions predominantly occurred in the posterior region of the mesoderm (Fig. 3I). Together, this tissue-level analysis indicates that the timing and posterior location of early cell internalisation events during *Tribolium* gastrulation correlate with enhanced tissue crowding associated with apical constriction during cell proliferation.

### Interpretable machine learning analysis associates local decrease in apical area and increase in nuclear crowding with out-of-plane divisions

To further investigate tissue properties that might affect division orientation, we used machine learning to ask how the local tissue environment relates to division orientation at the individual cell level (Fig. 4A). Focusing on the apical surface, we quantified a variety of features reflecting local tissue crowding and overall cell shape in live movies of gastrulating *Tribolium* embryos (Table S1). To enable comparisons between cells dividing at different times, we time-aligned the data for each individual cell, so that the last time-point prior to cell cleavage corresponds to *t* = 0 for each cell (Fig. 4A). Since the time resolution of our movies was 4 minutes, the frame at normalised *t* = 0 could capture cells at slightly different stages between late metaphase and anaphase. To reduce the effects of this asynchrony on our analysis, we averaged feature values within time bins spanning 2 acquisition points. This generated a dataset of features reflecting the apical geometry and local environment of each dividing cell over 7 time-points in the 30 min preceding cell division.

**Figure 4.**
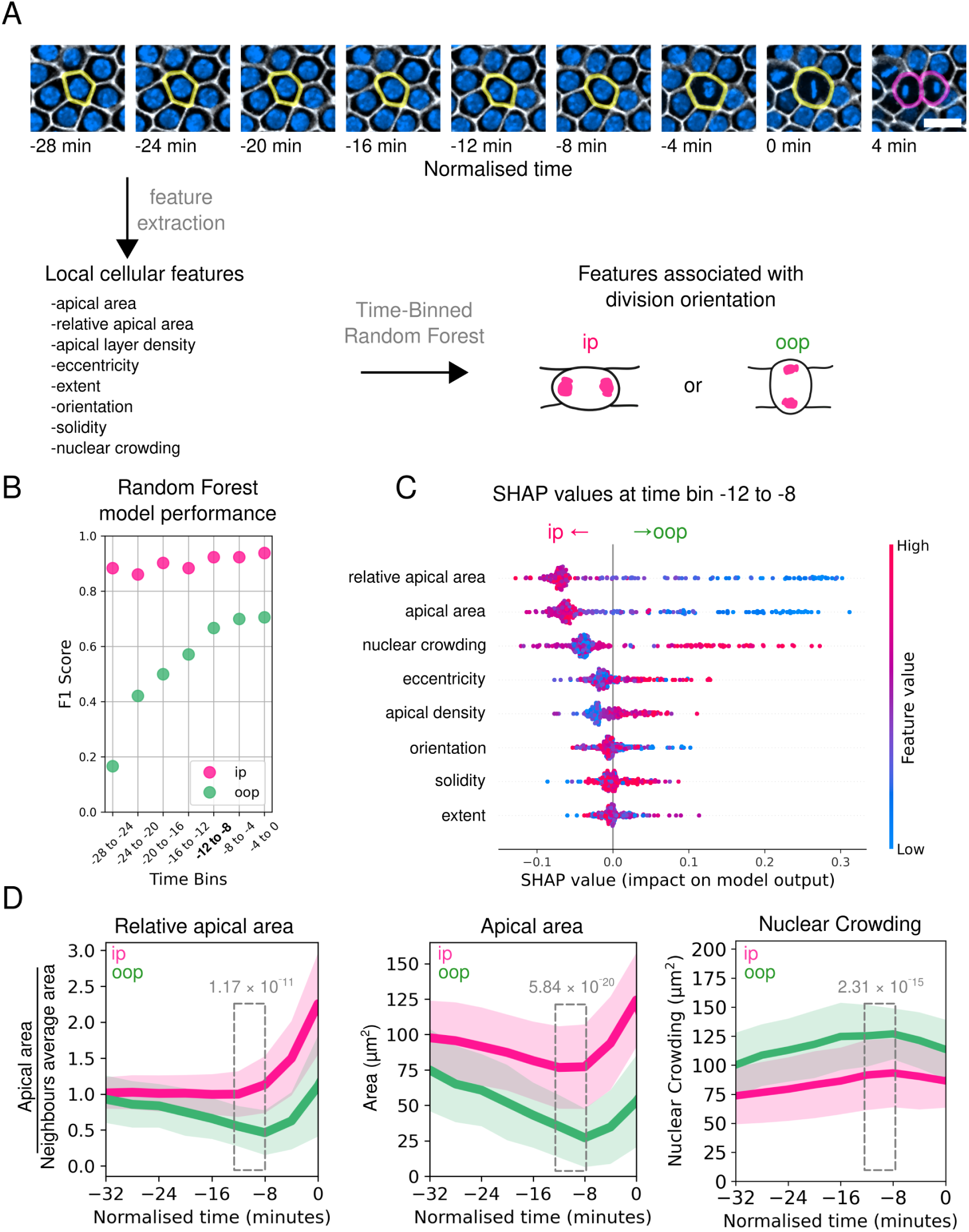
Machine-learning analysis identifies geometric predictors of division orientation in the *Tribolium* mesoderm. (A) Overview of the interpretable machine learning pipeline to establish the features most predictive of division orientation. Dividing cells were tracked over the 28 minutes preceding cell division. For each cell, *t* = 0 was defined as the last time-point preceding cell cleavage. For each dividing cell, features describing apical cell geometry and local tissue environment were extracted over time. A time-binned Random Forest Classifier, interpreted using SHAP values, was used to determine the features most predictive of division orientation. Scale bar, 10 µm. (B) Random Forest classifier performance in predicting cell division orientation from geometric and tissue density features. Feature values were binned across two successive time points over the 28 minutes preceding cell division. F1-scores (harmonic mean of precision and recall) for predicting in-plane (ip, magenta) and out-of-plane (oop, green) divisions are shown for successive time bins. (C) Summary plot of SHAP values for features measured in a time bin between 12 and 8 min prior to cell division. Features are ranked by their predictive contribution, with the most predictive feature at the top. Each point represents a single prediction of either in-plane (magenta) or out-of-plane (green) divisions. The position of the point on the x-axis shows how much that feature increases or decreases the likelihood of the prediction being in-plane or out-of-plane. Negative SHAP values indicate that the feature pushes the prediction toward in-plane, while positive values favour out-of-plane predictions. (D) Relative apical area (cell apical area relative to the average of neighbours within a 20 µm window), average apical cell area and nuclear crowding over time for cells dividing in plane (pink, *n* = 189) and out-of-plane (green, *n* = 61). Lines represent mean values calculated from cells tracked in *N* = 3 embryos; the shaded area represents the standard deviation. The Mann-Whitney p-value is reported for data averaged between 16 and 8 min prior to division.

We then identified which local cellular features most strongly associated with out-of-plane division orientation. We first trained a Random Forest Classifier^47^ on our dataset to predict whether a cell would divide in- or out-of-plane (Fig. 4A, see Methods for details). Assessing multiple time-points allowed us to track how the model’s predictive power evolved at different stages leading up to cell division. To assess model performance, we used the F1 score^48^, a measurement of the model performance that is robust to class imbalance between the frequent in-plane and rare out-of-plane divisions (see Methods). We found that while the classifier performed well for predicting in-plane divisions across all time-points, we observed a progressive increase in the predictive power for out-of-plane division, plateauing at time bin -12 to -8 min (Fig. 4B). This finding means our group of features is predictive of out-of-plane division orientation from 12 min before division onwards.

To determine which features were most predictive of division orientation, we used Shapley Additive Explanations (SHAP) values to analyse the model’s decision-making process at -12 to -8 min before division (Fig. 4C). SHAP analysis quantifies how much each feature contributes to a specific prediction of a machine learning classifier^49,50^. Unlike traditional feature importance scores, which only reflect a feature’s overall influence on model accuracy, SHAP values describe how the specific value of each feature affects individual predictions. This allowed us to not only identify the features that most strongly drive model performance, but also to uncover how individual feature values affect specific model predictions. SHAP analysis showed that the relative cell area, absolute cell area, and nuclear crowding features were the strongest drivers of model performance in the −12 to −8 min time window (Figs. 4C and S5A). Specifically, small apical cell area—both in absolute terms and relative to nearest neighbours—and high local nuclear crowding were the features most predictive of out-of-plane division orientation. Other features contributed less to distinguishing in-plane from out-of-plane divisions, although some still reflected aspects of crowding (e.g. higher apical density was also associated with out-of-plane divisions; Fig. 4C and S5A). This pattern is clear from the SHAP dependency plots: absolute area, relative area, and nuclear crowding display pronounced bimodal distributions, meaning their values have a strong impact on discriminating between in- and out-of-plane divisions, whereas the other features show progressively flatter distributions, indicating lower discriminative power (Fig. S5A). Plotting the three most predictive features over time revealed that cells dividing out-of-plane displayed smaller absolute and relative apical areas and experienced higher nuclear crowding compared with cells dividing in-plane throughout the 30 min preceding division (Fig. 4D). In contrast, features not directly related to crowding, such as solidity or eccentricity, did not display consistent trends (Fig. S5B).

Finally, we asked if cell extrusions also correlated with features associated with high local tissue crowding. The low frequency of cell extrusion events (*n* = 22 observed in 3 embryos) prevented effective use of our machine-learning SHAP analysis. However, we did observe that, similar to cells dividing out-of-plane, cells undergoing extrusion displayed smaller apical area and higher nuclear crowding compared to non-extruding mesodermal cells (Fig. S5C). This suggests that local tissue crowding may also drive cell extrusion. Together, our systematic machine-learning analysis combined with dynamic analysis of features describing cell shape and local environment strongly suggest that, in the *Tribolium* mesoderm, out-of-plane divisions and cell extrusions are driven by local tissue crowding.

### Perturbation experiments suggest early cell internalisation results from apical constrictions combined with cell proliferation

Our results show a strong correlation between increased tissue crowding in the apically constricting *Tribolium* mesoderm, and out-of-plane division and cell extrusion events. We thus asked whether abrogating apical constrictions by inhibiting mesoderm specification would decrease the occurrence of these early internalisation events. We performed RNAi mediated knock down of *Tc-twi*, a key driver of mesoderm specification and of enhanced apical contractility in the mesodermal region^30^. We confirmed that the ventral tissue in *Tc-twi* knockdown (*Tc-twi-KD*) embryos failed to undergo folding (Fig. 5A), and that cells did not display apical constriction (Fig. 5B). Cells in *Tc-twi-*KD embryos still divided, with the average apical area of daughter cells roughly half that of the mothers (Fig. 5B). However, the overall ventral tissue area showed a smaller reduction in *Tc-twi-KD* embryos compared to control embryos (Fig. 5C), in line with the abrogation of apical constriction. Manual tracking showed that *Tc-twi-KD* embryos displayed no out-of-plane divisions or cell extrusion events (Video S3, *N* = 3 embryos). These observations support the hypothesis that both out-of-plane divisions and extrusions events are a consequence of increased tissue crowding resulting from apical constrictions.

**Figure 5.**
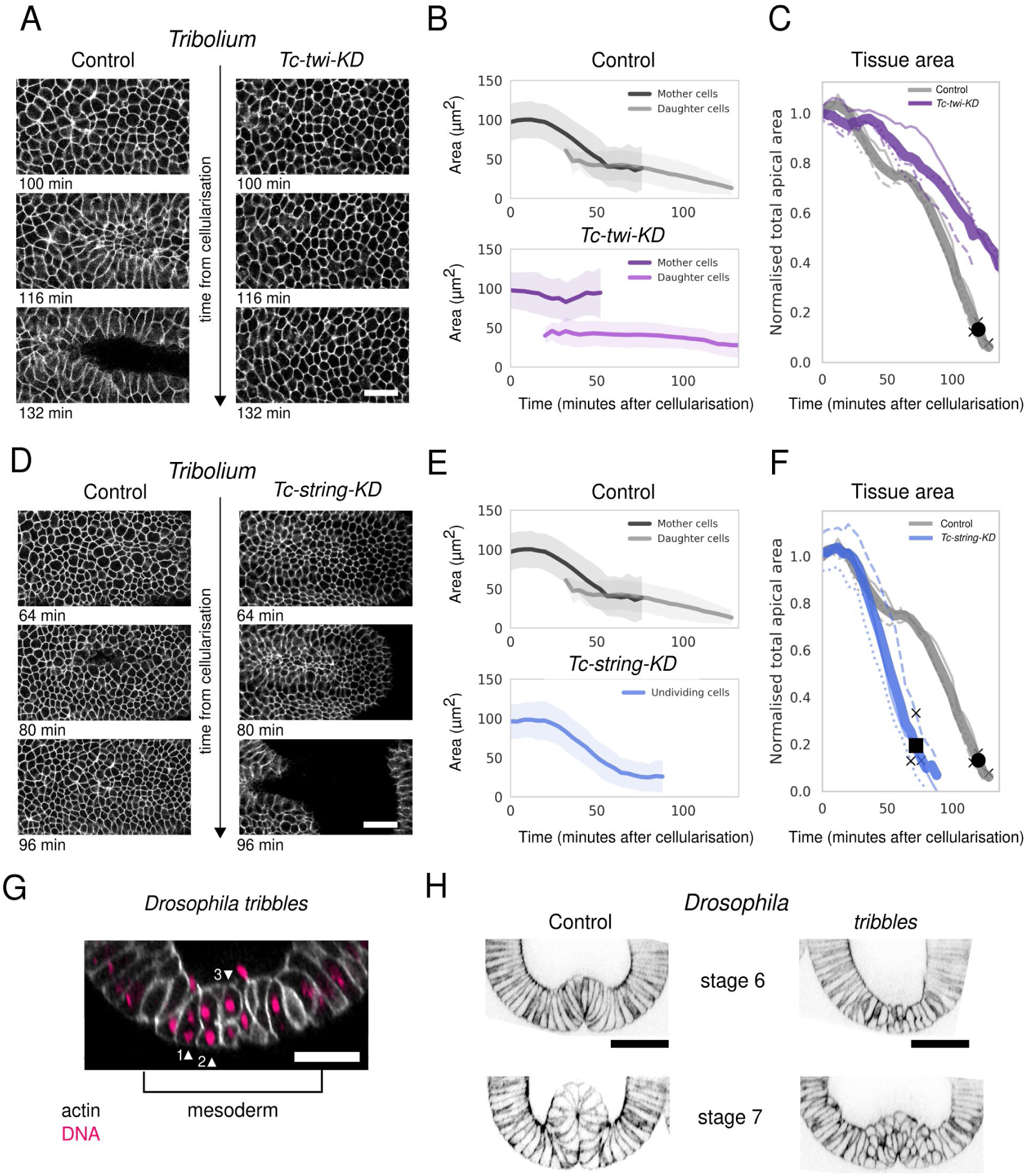
Perturbation experiments suggest that proliferation in an apically contracting mesoderm leads to out-of-plane divisions. (A) Representative time-lapse of the apical surface (average projection of grazing section of 4∼µm) of a control *Tribolium* embryo (left) and an embryo injected with *Tc-twi* dsRNA (right). The *Tc-twi-KD* embryo does not display apical contraction. Scale bar, 40 µm. (B) Apical cell area over time of the ventral region in control (top) and *Tc-twi-KD* (bottom) embryos, for cells before division (mother cells) and after division (daughter cells). Sample sizes: control mothers (*n* = 170), control daughters (*n* = 235), *Tc-twi-KD* mothers (*n* = 150), *Tc-twi-KD* daughters (*n* = 222) in *N* = 3 embryos per condition. As cell area transiently increases during mitotic rounding, cell area in the 8 minutes preceding division was excluded from mother cell tracks. Solid line: mean, shaded area: ± standard deviation. (C) Normalised sum of the apical area of all tracked cells over time in control and *Tc-twi-KD* embryos, used as a proxy for relative tissue area. For each embryo, total apical area was normalised to its initial value. Dashed lines: individual embryos (*N* = 3 per condition), thick line: mean. Crosses mark furrowing times for individual embryos. The black circle indicates the mean invagination times for control (121 min). (D) Representative time-lapse of the apical surface (average projection of grazing section of 4∼µm) of a control embryo (left) and an embryo injected with *Tc-string* dsRNA (right). The *Tc-string-KD* embryo displayed apical constrictions and forms a ventral furrow earlier than control. Scale bar, 40 µm. (E) Cell apical area over time in control (top) and *Tc-string-KD* (bottom) embryos, for cells before division (mother cells) and after division (daughter cells). Sample sizes: control mothers (*n* = 170), control daughters (*n* = 235), *Tc-string-KD* undivided cells (*n* = 99) in *N* = 3 embryos per condition. As cell area transiently increases during mitotic rounding, cell area in the 8 minutes preceding division was excluded from mother cell tracks. Solid line: mean, shaded area: ± standard deviation. The top panel shows the same data as in top panel B. (F) Normalised sum of the apical area of all tracked cells over time in control and *Tc-string-KD* embryos, used as a proxy for relative tissue area. For each embryo, total apical area was normalised to its initial value. Dashed lines: individual embryos (*N* = 3 per condition), thick line: mean. Crosses mark furrowing times for individual embryos. The black circle and black square indicate the mean invagination times for control (121 min) and *Tc-string-KD* (72 min) embryos, respectively. The control values are the same data as in panel C. (G) Cross-section of a representative *Drosophila tribbles* mutant embryo stained with DAPI to mark the DNA (magenta) and phalloidin to mark cell outlines (greyscale). Arrowheads 1,2 and 3 highlight cells at different stages of cell division in three neighbouring mesodermal cells, highlighting out-of-plane division orientation. Scale bar, 20 µm. (H) Representative images of control and *tribbles Drosophila* embryos at two successive stages of mesoderm invagination. Embryos were stained for phalloidin (inverted contrast is displayed). The *tribbles* embryo displays delayed furrow invagination and early internalised cells compared to control. Scale bar, 40 µm.

As reduction in apical constriction and tissue crowding abrogated cell internalisation events, we asked whether decreasing tissue crowding by blocking cell divisions would have the same effect. We thus blocked cell divisions by knocking down the *Tribolium* homolog of Cdc25 (*Tc-string*, *string* in *Drosophila*), a phosphatase shown to be required for post-blastoderm mitotic entry in *Drosophil*a^51–53^. We optimised the time of dsRNA injection to target the divisions occurring during gastrulation without disrupting previous cell division rounds (see Methods). Mesoderm cells of *Tc-string*-KD embryos did not display any divisions but still displayed apical constrictions (Fig. 5D,E), and we did not observe any cell extrusions in the mesoderm of *Tc-string*-KD embryos (Video S4). Interestingly, we further observed that ventral furrow invagination occurred ∼50 min earlier in *Tc-string-KD* embryos compared to controls (Fig. 5F). Together, these perturbation experiments suggest that early cell internalisation through out-of-plane division or extrusion in the *Tribolium* mesoderm occurs due to tissue crowding resulting from both apical constrictions and cell divisions. Furthermore, blocking cell division considerably accelerated the rate of gastrulation in *Tribolium*.

### Premature mesodermal mitotic entry in the *Drosophila tribbles* mutant results in out-of-plane divisions

Since cell division affects cell internalisation timing in *Tribolium*, we decided to investigate the effect of cell divisions in the *Drosophila* mesoderm, which undergoes apical constriction but does not normally display cell divisions^34,36^. In *Drosophila*, ventral cell divisions during gastrulation are inhibited by *tribbles*, which acts to degrade String^35^. Consistently, in *Tribolium Tc-tribbles* expression is confined to the extraembryonic serosa, where cell divisions do not occur (Fig. S6A). We thus analysed cell behaviour in *Drosophila tribbles* mutant embryos, which display premature cell divisions in the mesoderm, disrupting apical constrictions and ventral furrow formation^54^. As hypothesised from our observations in *Tribolium*, mitosis during mesoderm invagination in *Drosophila* resulted in out-of-plane divisions in the mesoderm (Fig. 5G). Furthermore, *tribbles* embryos formed a ventral furrow later than in control embryos (Fig. 5H, Fig. S6B), mirroring the delayed mesoderm invagination in *Tribolium* wild-type embryos compared to *Tc-string-KD* embryos. Together, these observations suggest that cell divisions during gastrulation hinder tissue invagination, and that out-of-plane divisions may represent a conserved cellular response to this mechanical challenge (Fig. 6).

**Figure 6.**
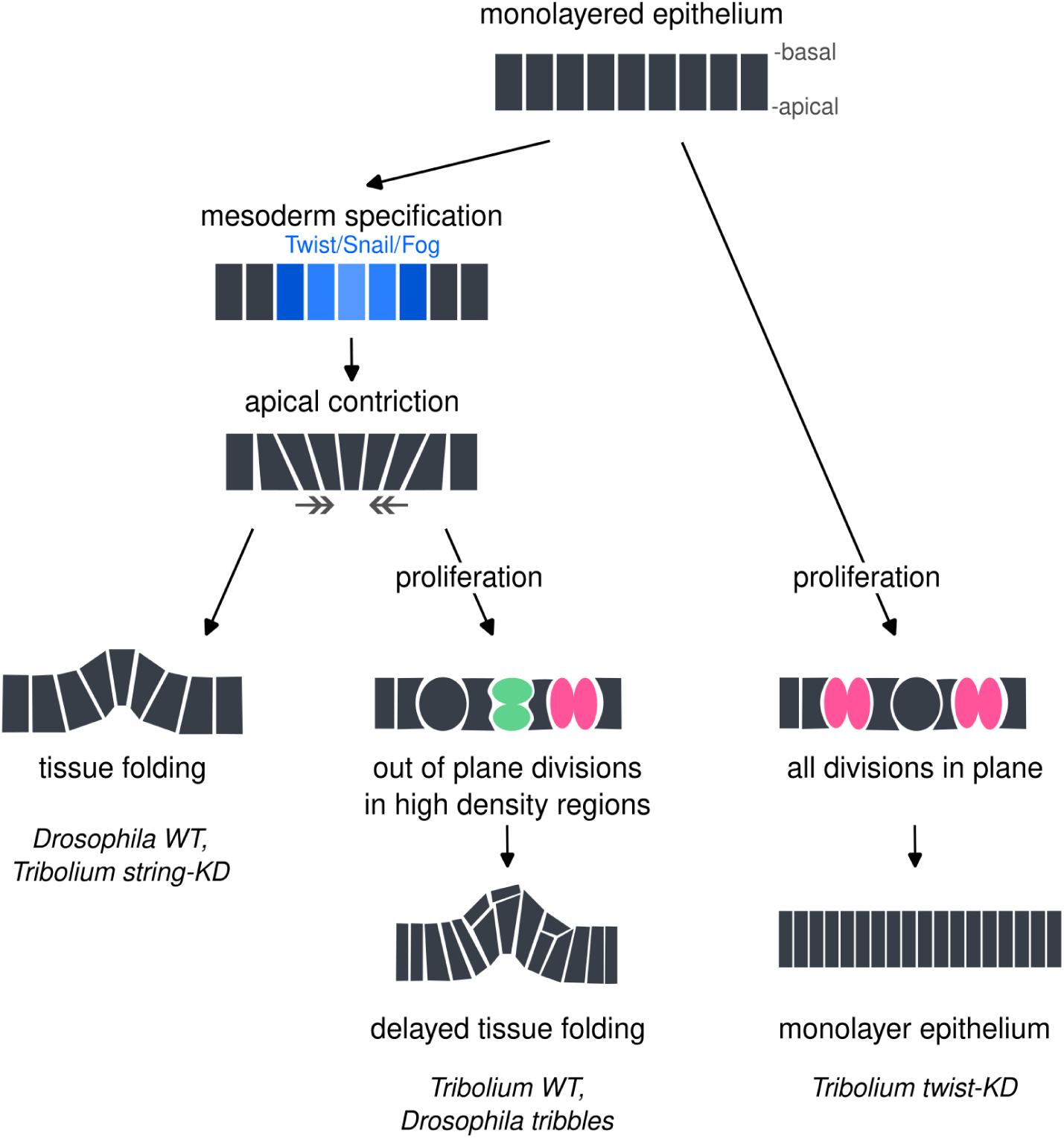
Distinct cellular behaviours during mesoderm invagination. Schematic of the different cellular behaviours contributing to tissue internalisation during insect gastrulation. In both *Drosophila* and *Tribolium*, mesoderm specification leads to apical constriction of the mesoderm cells, which by itself can drive tissue folding (left). When apical contractions are coupled to cell proliferation, tissue folding is delayed and out-of-plane divisions lead to early cell internalisation (middle). Proliferation without apical constrictions does not result in out-of-plane divisions (right), suggesting that apical contraction contributes to tissue crowding leading to out-of-plane division.

## DISCUSSION

By embedding cell shapes into a common low-dimensional space, the morphospace constitutes a quantitative framework for comparison of morphogenetic behaviours between conditions and across species. For instance, the mesoderm morphospace trajectory identified in *Drosophila* (Fig. 1C, E) provides a quantitative reference to assess how perturbations affect cell shape dynamics during gastrulation. More generally, morphospace analysis enables quantification of cell shape dynamics and variability, and the identification of rare or divergent morphological behaviours. Using morphospace analysis, we show here that despite similar tissue-scale internalisation, *Tribolium* and *Drosophila* mesoderm cells follow markedly different cell shape trajectories (Fig. 1). Focusing on time-points of highest shape variability in morphospace, we found that a subset of *Tribolium* mesodermal cells internalises prior to tissue folding via out-of-plane divisions and, to a smaller extent, cell extrusions (Figs. 2,3).

Cell division during gastrulation is observed across the animal tree, but examples of out-of-plane division resulting in one daughter cell leaving the epithelium are rare. In several crustaceans^55–57^, and the cephalochordate *Amphioxus lanceolotum*^58^, cell division contributes to folding the epithelial plane during gastrulation, but both daughter cells remain in the epithelium. In the mouse primitive streak, non-apical mitosis has been proposed to facilitate EMT-driven cell delamination^59,60^, but again, both daughter cells initially remain within the epithelium. In contrast, out-of-plan divisions, like we observe in *Tribolium*, have been described in several cnidarian species during gastrulation^61–65^. Apart from cnidarians, out-of-plane divisions have not, to our knowledge, been reported as a direct driver of mesoderm internalisation.

In contrast, many animals repress^66–68,12,69^ or finely tune cell division in mesoderm/endoderm during gastrulation. For instance, in the frog *Xenopus laevis*, division is repressed in some cells but required in others for gastrulation to proceed^70^. The consequences of inappropriate division during gastrulation have been investigated in *Drosophila*, where mitotic rounding in *tribbles* mutants antagonises apical constriction during ventral furrow formation by reducing apical RhoA-mediated myosin contractility, and disrupts coordinated tissue constriction^54^. Consistent with this, our data show that abrogating cell division in the *Tribolium* mesoderm accelerates tissue folding, while inducing cell division in the *Drosophila* mesoderm delays folding (Fig. 5A-F). Together, these observations suggest that cell proliferation and apical contractility are antagonistic processes during epithelial tissue folding in these embryos (Fig. 6). Our work suggests that the induction of *tribbles* in the mesoderm to repress divisions, and thereby accelerate gastrulation, could have been a key evolutionary step towards the rapid embryonic development exhibited by *Drosophila*^71–73^.

Our machine learning-assisted analysis showed that out-of-plane divisions specifically associate with locally increased tissue crowding. Furthermore, when apical contractions were abrogated, all cells divided within the epithelial plane, suggesting that apical constriction is a driver of tissue crowding. Crowding has been shown to induce out-of-plane divisions in the mammalian skin, where cells will divide in plane until they reach a density threshold above which they reorient mitotic spindles perpendicular to the epithelium^45^. Studies in epithelial cultures have further demonstrated that compressing an epithelial monolayer can cause cells to reorient the spindle perpendicular to the plane of epithelium^46^. It will be interesting to investigate which molecular mechanisms lead to spindle re-orientation during out-of-plane divisions in the *Tribolium* mesoderm.

Out-of-plane divisions have recently been proposed to provide a mechanism to buffer mechanical compression forces in the cephalic region in *Drosophila* and other dipterans^74,75^. It is tempting to speculate that out-of-plane divisions play a similar role in the *Tribolium* mesoderm, and in the mesoderm of *Drosophila tribbles* mutants, releasing tissue pressure build-up upon tissue crowding that is induced by the combined effect of apical constriction and cell proliferation. This suggests that dividing out-of-plane in a constricting tissue may constitute a conserved response to the mechanical constraints imposed during gastrulation^76,77^. Cell proliferation during gastrulation has been reported in many insects beyond *Tribolium*^78–82^, but whether any exhibit crowding-induced out-of-plane divisions requires future work. Quantitative investigation of the relationship between proliferation, apical constriction and division orientation across species is now needed to reveal both the evolutionary origins and the mechanical principles underlying the cellular behaviours driving gastrulation.

## RESOURCE AVAILABILITY

### Lead contact

All plasmids generated in this study are available from the lead contact Ewa K. Paluch upon request. The datasets generated during this study will be deposited in the BioImage Archive prior to publication, and all custom analysis code are available from the corresponding authors.

## ACKNOWLEDGMENTS

We thank Jee In Kim for help with preliminary data analysis. We thank Gergely Flamich for support with the machine learning analysis. We are grateful to Michael Akam and Emilia Santos for providing laboratory space for M.B. We thank Alba Diz-Muñoz, Nicoletta Petridou, and the Advanced Facility for Light Microscopy (AFLM, Heidelberg) for hosting and support with live imaging. We acknowledge the Cambridge Advanced Imaging Centre (CAIC) for support with image acquisition. We thank Augusto Borges, Erik Clark, Anna Foix Romero, Bénédicte Sanson, Marta Urbanska, and Iskra Yanakieva for critical comments on the manuscript. We also thank Yohanns Bellaïche, Siegfried Roth, Guillaume Salbreux, Tasmin Sarkany, Daniel St Johnston, Quentin Vagne, and other members of our research communities for helpful discussions and feedback throughout the project.

M.B. was supported by the Wellcome Trust Developmental Mechanisms MPhil + PhD Programme. This work was supported by the European Research Council (ERC consolidator grant *NanoMechShape* to E.K.P.), by a Leverhulme Trust Prize awarded to E.K.P., and a Company of Biologists travel fellowship to M.B.. W.P. acknowledges financial support from the Herchel Smith Fund through a Herchel Smith Postdoctoral Fellowship. M.A.B was supported by the Deutsche Forschungsgemeinschaft (Research Fellowship), the Isaac Newton Trust (Research Grant), and the Department of Zoology, University of Cambridge.

## AUTHOR CONTRIBUTIONS

M.B., E.K.P., and M.A.B. designed the study. M.B. and M.A.B performed the experiments. M.B. carried out data analysis, developed computational pipelines, generated figures, and wrote the original manuscript draft. W.P. developed the morphospace analysis framework and contributed to data analysis. E.K.P. and M.A.B. supervised the research, contributed to experimental design and interpretation of the results. M.B, M.A.B. and E.K.P. wrote the paper. All authors discussed the results.

## METHODS

### ANIMAL HUSBANDRY

Oregon R. *Drosophila melanogaster* fly stocks were maintained at 25°C on standard recipe food and were transferred every 2-3 weeks. Eggs were collected following standard procedures by placing flies in cylindrical plastic cages, with a mesh on one end to ensure air circulation, and a removable apple agar plate on the other side. The plates were streaked with yeast paste to stimulate egg laying. Flies carrying the loss of function *tribbles^EP^*^1119^ allele^35^ were a gift from Bénédicte Sanson. They were balanced over a TM3 balancer with a hunchback-lacZ marker (BDSC:78357), so that homozygous *tribbles^-/-^* could be recognised by the absence of signal when staining against *lacZ* transcript.

*Tribolium castaneum* beetles were reared at 30°C in plastic boxes equipped with mesh-covered windows in the lid to ensure proper air-flow. Beetles were maintained on a diet of coarse flour, which was prepared by mixing 1 kg of Organic Whole Wheat Flour (Prior’s Flour, U.K.), approximately 50 g of dry yeast powder (Sainsbury’s supermarket), and 1 g of Fumagilin-B (Medivet #12415). All components of the diet were sieved through a 700 μm steel sieve (Retsch test sieve 200mm x 50mm) to remove large particles before mixing. Food was replaced every 1-2 weeks. Once a month, an overnight egg lay was collected and aged for one month at 30°C to produce the next generation of beetles. For egg collection, beetles were separated from the coarse flour mix using an 800 μm steel sieve (Retsch test sieve 200 mm x 50 mm) and transferred to Prior’s Flour organic white flour, that had been previously sieved using a 300 μm steel sieve (Retsch test sieve 200 mm x 50 mm). Eggs were subsequently collected from the white flour using a 300 μm steel sieve. Incubators were maintained at a relative humidity between 40-60%. For more comprehensive details on *Tribolium* husbandry, refer to “The Beetle Book”^83^.

To our knowledge, the gender of embryos has not been reported as having an effect on gastrulation. We therefore did not record it in our studies.

### EMBRYO DECHORIONATION AND FIXATION

*Drosophila* flies were allowed to lay eggs in continuous 20-minute intervals. Embryos were fixed at specific time points, namely 180, 200, 220, and 240 minutes after egg laying. Prior to fixing, embryos were dechorionated for 3 minutes in a small mesh basket (with a mesh aperture of 250 μm) using a bleach solution (Thermo Scientific Chemicals 10527752) diluted with ddH_2_O to achieve a final 2.5% hypochlorite concentration, following standard procedures^84^. Embryos were then washed several times with a squirt bottle containing double distilled water to remove any bleach trace. The basket was placed on a paper tissue to absorb most of the water, and the embryos which rested at the bottom of the basket were transferred to a tube containing heptane using a fine brush. Embryos were fixed on a rocker in a 1:1 solution of 37% paraformaldehyde and heptane, for 5 minutes at room temperature. Depending on the number of embryos, we fixed them either in a 1.5 ml plastic eppendorf tube, or in a 5 ml glass vial. Embryos were then washed twice in 1x PBS (137 mM NaCl, 2.7 mM KCl, 10mM Na_2_HPO_4_, 1.8 mM KH_2_PO_4_) to remove the fixative. PBS was then removed and replaced with clean heptane. Usually at this stage, the vitelline envelope surrounding the embryo is removed using MeOH, which allows for the penetration of antibodies and other staining molecules in the embryonic tissues. However, MeOH destroys the epitope recognised by phalloidin. To allow for phalloidin staining, we therefore followed a protocol to remove the vitelline envelope without the use of MeOH^84^. The embryos in clean heptane were sucked into a glass pipette, whose neck was cut with forceps and sanded to create an aperture of 2-3 mm in diameter. Embryos were then dropped in the centre of a circular Whatman filter paper No. 3 (Camlab #1003-070). Heptane was allowed to evaporate from the paper for less than 30 seconds, as drying the embryos longer can damage the samples. The embryos were transferred to a double-stick tape placed at the bottom of a 100 mm petri dish by delicately placing the filter paper into the dish with the side where the embryos are attached facing downwards. The embryos were immediately covered in PBS, and the vitelline envelope was manually removed. To do so, a 3-4cm segment of Tungsten 0.375 mm wire (Thermo Fisher #043995.H1) was cut, heated over a flame and sharpened by striking it onto a Sodium Nitrite stick (McCRONE #107-SK). The sharpened wire segment was inserted into a 0.8mm needle (Terumo Agani #8AN2150R1) which was then glued onto a p1000 tip by melting the plastic with a flame and quickly bringing the melted pipette tip and wire together. This tool was then used to carefully break the vitelline envelope of the fixed embryos and dissect the embryos out. After dissecting approximately 200 embryos, they were transferred to a fresh tube containing PBS using a 200 μm pipette with a plastic pipette tip briefly pre-washed with 1% Bovine Serum Albumin (BSA) in phosphate buffered saline (PBS) solution to prevent the embryos from sticking to the plastic. The pipette tip was cut using a razor blade to create an aperture of 2-3 mm.

*Tribolium* beetles were allowed to lay eggs for one hour at 30°C, as their development takes an average of three times longer than *Drosophila*. Beetles were then sieved out, and the white flour containing the eggs was allowed to age at 25°C for 18, 20, 22, 24, 26, and 28 hours. Collected embryos were initially placed in small mesh baskets (with a mesh aperture of 250 μm) and underwent a series of rinses in type 2 water to eliminate any residual flour. This was achieved by placing the basket containing the embryos in a 50 mm petri-dish filled with water for a 30-second rinse, during which the liquid was drawn up from outside the basket and then squirted onto it with a plastic 3 ml pipette. Subsequently, the basket was moved to a petri-dish containing Sodium hypochlorite (6-14% active chlorine, Merk #7681-52-9) diluted with water to achieve a final 2.5-5% hypochlorite concentration. The embryos were rinsed in it for 3 minutes with a plastic 3 ml pipette to dechorionate them. The embryos were then thoroughly rinsed by passing them sequentially in 6 petri-dishes filled with water. Finally, the dechorionated embryos were transferred out of the basket into a 1.5 mL fresh tube containing heptane with a fine brush. Dechorionated embryos float in water, so they were easily separated from the non-dechorionated ones. The embryos were fixed and devitellinised as described above for *Drosophila*, with the difference that *Tribolium* embryos were fixed for 7.5 minutes instead of 5 minutes. Longer fixation times of 10 minutes were sometimes needed for the most delicate blastoderm stages.

### EMBRYO STAINING

To visualise RNA transcripts in *Drosophila* embryos, we performed in situ Hybridisation Chain Reaction (HCR) v3.0^85^ using probes and hairpins produced by Molecular Instruments, following the protocol for whole-mount fruit fly embryos (https://files.molecularinstruments.com/MI-Protocol-HCRv3-FruitFly-Rev6.pdf). The protocol was modified according to published literature^86^: fixed embryos were not treated with ethanol, xylene, or proteinase K prior to staining, and, to reduce viscosity and allow the embryos to settle more easily at the bottom of the tube, the percentage of dextran sulphate in the probe hybridisation and amplification stock buffers was reduced from 10% weight per volume (w/v) to 5% w/v. Additionally, the probe hybridisation, amplification and wash solution were further diluted to 40% with water to allow phalloidin staining. The mRNA sequences used to produce the HCR probes are identified by the following FlyBase gene and isoform IDs: FBtr0080739 for *snail* (isoform *sna-RA*), and FBtr0100130 for *twist* (isoform *twi-RB*). To stain against the lacZ transcripts we designed probes against the sequence NCBI: NC_000913.3: c366305-363231. For each targeted gene, 20 pairs of short probes were ordered, as recommended by Molecular Instruments. Following HCR, embryos were incubated for 1 hour with 1 ng/μl DAPI (ThermoFisher Scientific) and in 5 ng/μl of DyLight650 Phalloidin (Cell Signaling T567) at room temperature to stain nuclei and filamentous actin. Embryos were then washed 3 times in 5x SSC + 0.1% Triton X-100 (5x SSCT). Embryos were imaged shortly after staining, as the phalloidin signal degrades significantly within a week.

HCR staining in *Tribolium* embryos was performed following the same procedure as described above for *Drosophila*, with the exception that the concentration of HCR probes and hairpins was doubled during the hybridisation step to enhance the signal (final concentration: 4 nM/mL)^87^. The mRNA sequences used to produce HCR probes against mesodermal gene transcripts can be found in the iBeetle database (https://ibeetle-base.uni-goettingen.de/) by searching for the following gene identifiers: TC032125 for *Tc-sna*, TC014598 for *Tc-twi*, TC033074 for *Tc-tribbles*, and TC001885 for *Tc-string*. DAPI and phalloidin staining following HCR was performed as described above.

For tubulin antibody stainings in *Tribolium*, fixed embryos were first treated with a blocking solution (5% Normal Goat Serum and 0.1% BSA in 1x PBT) for 30 minutes at room temperature with gentle agitation using a sample rack attached to a nutator. Embryos were then incubated with the primary antibody mouse monoclonal anti-acetylated-α-Tubulin (Sigma #T6793) at a 1:100 dilution and incubated overnight at 4°C with gentle agitation. The embryos were then subjected to four 15-minute washes in 1x PBT and 0.1% Triton X-100 at room temperature by placing the tube containing them horizontally on the nutator. This was followed by a 30-minute incubation in the blocking solution at room temperature. The embryos were then incubated for 2 hours at room temperature with goat anti-rabbit Alexa Fluor 555 (Invitrogen #A-21429) secondary antibody diluted 1:200, in a 1:1 mixture with 100% glycerol for storage. This step was followed by four 15-minute washes and a final 30-minute wash in 1x PBT at room temperature, with agitation throughout. DAPI and phalloidin staining following HCR was performed as described above.

In each experiment, embryos were stained for transcripts or antibodies using a combination of Alexa Fluor 488, Alexa Fluor 546, Alexa Fluor 555, and/or Alexa Fluor 594.

### IMAGE ACQUISITION FOR FIXED SAMPLES

Embryos were mounted immediately after staining in SlowFade Diamond Antifade Mountant (ThermoFisher # S36972) on glass microscope slides (Thermo Scientific) with 1.5 coverslips (Corning #2850-22). For *Drosophila* embryos, 1.5 coverslips were also cut and used as bridges to prevent embryos from being squashed. *Tribolium* embryos were mounted using a small amount of Plasticine at the 4 corners of the coverslip, gently pushed down with forceps until the sample touched it. Clear nail varnish was brushed along the coverslip perimeter to seal the edges of the slide.

Microscopy was performed at the Department of Zoology Imaging Facility (University of Cambridge) on an Olympus FV3000 confocal microscope using an Olympus UPlanSApo 30x 1.05 NA silicon immersion oil objective using a physical pixel size of 0.23 μm x 0.23 μm, and a Z-stack step size of 0.79 μm. Acquired images were 12-bit with a 1024x768 scan format and a 2 μs/pixel dwell time. The number of slices varied with the thickness of the sample. For the acquisition of the images used to extract 3D cell shapes, *Drosophila* embryos were aligned to capture the mesodermal tissues just posterior to the cephalic furrow, while *Tribolium* embryos were aligned based on the *Tc-eve* signal to capture the first three parasegments (Patel et al. 1994). All imaging channels were acquired sequentially to minimise crosstalk. The laser lines and collection windows were: 405 laser and 443-472 nm window for DAPI; 488 laser and 500-536 nm window for Alexa Fluor 488; 561 laser and 566-584 nm window for Alexa Fluor 546 and Alexa Fluor 555; 594 laser and 610-631 nm window for Alexa Fluor 594; 640 laser and 663-713 nm window for DyLight650.

### DESIGN OF mRNA TEMPLATES FOR MICROINJECTION IN *TRIBOLIUM*

To visualise membranes and nuclei in live imaging experiments in *Tribolium*, we designed two fluorescent reporters coding for a signal peptide that localises either to the membranes or the nuclei, linked to a fluorescent protein. The plasma membrane was labelled using a fluorescent protein fused to the N-terminal palmitoylation sequence of GAP43, which directs the protein to the plasma membrane. This is a small protein that localises to the inner surface of the plasma membrane and has been successfully used as a membrane marker in *Tribolium*^37^. To mark the nuclei, the *Tribolium v*ariant histone H2A (H2Av) sequence was identified by aligning the *Drosophila* H2Av sequence to the *Tribolium* genome using the blastn suite (NIH). We fused these markers to bright fast folding fluorescent proteins: the yellow-green fluorescent protein mNeonGreen^88^ for Gap43, and the red fluorescent protein mScarlet-I^89^ for H2Av. To enhance marker translation, upstream of the signal peptide we added intervening sequence (IVS) from *Drosophila* myosin heavy chain to facilitate mRNA export to the cytoplasm^90^, and the synthetic AT-rich 21 bp sequence (Syn21) to promote translational initiation^91,92^. Downstream of the fluorescent reporter sequence we added the highly efficient baculovirus p10 polyadenylation (polyA) signal^93,92^. The fluorescent constructs were synthesised and inserted in a pUC57 vector by Genescript (U.K.); the sequences were codon optimised for insects prior to synthesis. Once received, plasmids were transformed into DH5α competent cells (Zymo Research T3009) following standard protocols and mini-prepped using the Monarch Plasmid Miniprep Kit (#T1010S). The obtained plasmids are identified as *pUC57[G43-mNeonGreen]* and *pUC57[Tc-H2Av-mScarlet-I]* and are available upon request.

### CAPPED, SINGLE STRANDED RNA SYNTHESIS

To transiently label membrane, nuclei or non-muscle myosin II for live imaging in *Tribolium* embryos, we synthesised capped mRNA from three DNA plasmids: *pUC57[G43-mNeonGreen]* to mark the membranes (see above), *pUC57[Tc-H2Av-mScarlet-I]* to mark the nuclei (see above), and *pCS2+[Tc-sqh-GFP]* to mark non-muscle myosin II regulatory chain with Green Fluorescent Protein (kindly provided by Pavel Tomancak)^94^.

Constructs were amplified from the plasmids using a 5’-primer with an overhang containing the SP6 RNA polymerase gene core promoter sequence (ATTTAGGTGACACTATAG)^95^. PCR reactions were performed using either the Phusion High-Fidelity system (New England Biolabs #E0553L) or the Q5 High-Fidelity DNA Polymerase (New England Biolabs #M0491).

Amplicons generated from these plasmids were purified using the Monarch Spin gDNA Extraction kit (New England Biolabs #T3010S). Subsequently, they were precipitated and resuspended in nuclease-free water to achieve a concentration of approximately 100 ng/μl. Capped mRNA was then synthesised from the purified DNA amplicons using the SP6 mMESSAGE mMACHINE kit (Invitrogen #AM1340), following the manufacturer’s instructions. After digestion with TURBO-DNase, RNA was purified through a phenol:chloroform and chloroform extraction, followed by the addition of an equal volume of isopropanol for precipitation. RNA was stored in isopropanol at -20°C. Before the injection, RNA was collected via a 20 minute centrifugation at 13,000 rpm at 4°C, and resuspended in 5 μl of nuclease-free water. RNA concentration was quantified using a Nanodrop spectrophotometer.

### DOUBLE STRANDED RNA SYNTHESIS FOR GENE KNOCK-DOWNS

cDNA was extracted using the TRIzol reagent (Invitrogen #15596-026) from 0-72 hour old embryos and synthesised using the Superscript III First-strand Synthesis System for RT-PCR kit (Invitrogen #18080-051). The sequences of the *Tc-twi* gene (TC014598) and the *Tc-string* gene (TC001885) were amplified from the extracted cDNA using the following primers: *Tc-twi*-F GCTGATGGACCTGACCAACT, *Tc-twi-R* GCTTGAATTTCGGTGGAAAA; *Tc-string-F* TTTCCGATATTGAGCCCTTG, *Tc-string-R* TCCAGGTTTTGGATTTGAGC. PCR products were ligated into the plasmid pGEM-T Easy (Promega #A1360) or pCR4-TOPO (ThermoFisher #K450030), generating *pGEM[Tc-twist]* and *pCR4[Tc-string]*.

DNA was amplified from *pGEM[Tc-twi]* and *pCR4[Tc-string]* using M13 primers with T7 overhangs: *M13-18mer-F* TGTAAAACGACGGCCAGT, *M13-T7-R* GGGTAATACGACTCACTATAGGGCAGGAAACAGCTATGACC. PCR products were purified using the Monarch Spin gDNA Extraction kit and diluted to 1 µg/µl. Double stranded RNA (dsRNA) was generated using the MEGAscript T7 Transcription Kit (Invitrogen #AM1334), following the manufacturer’s instructions. The reaction was incubated at 37°C for a total of 4 hours. The RNA mixture was then incubated at 75°C for 5 minutes, after which it was left on the bench to cool naturally to room temperature, allowing the RNA to anneal and form double-stranded RNA. Subsequently, phenol-chloroform extraction was carried out. The sample was stored overnight at -20°C after addition of 0.1 volumes of Ammonium Acetate STOP Solution from the kit, and 2.5 volumes of Denatured Ethanol 99.5% (APC Pure #GWN9065-E). The next day, the sample was centrifuged at maximum speed at 4°C for 30 minutes. After removing the supernatant, 1 mL of fresh ice-cold 70% ethanol solution was added. The sample was centrifuged again at maximum speed for 30 minutes. The liquid was removed, and the pellet was allowed to air dry for 10 minutes. Finally, the RNA pellet was resuspended in 20 µL of water.

### *TRIBOLIUM* EMBRYO MICROINJECTION

Embryonic microinjection was performed following published procedures^5^. Adult beetles were placed in sieved Organic Plain Flour (Prior’s Flour) and allowed to lay eggs for 1 hour at 32°C. Adults were then sieved out of the flour, and the embryos were allowed to age for at least one hour. Eggs were dechorionated as described above for the collection of fixed embryos, except a weaker bleach solution was used (0.5% hypochlorite concentration) to be gentler. Embryos were washed in bleach solution for two rounds of 30 seconds, with a 30 second pause in between in ddH_2_O. Dechorionated embryos were transferred using a glass pipette (whose neck was cut with forceps and sanded to create an aperture of 2-3 mm in diameter) to a drop of water onto a 50x22 mm coverslip, held in place on a glass slide by a small amount of Plasticine.

The embryos were aligned on the coverslip with the anterior end of the embryo facing outwards using a single-strand paintbrush, or alternatively an eyebrow hair attached to a glass pipette with blue tack. To avoid suffocation, embryos were spaced to not touch each other. Once around 50 embryos were aligned, and the water evaporated, they were covered in a halocarbon oil mix, made up of 50% Halocarbon oil 700 (Sigma Aldrich #H8898-100ML) and 50% Halocarbon 27 (Sigma Aldrich #H8773-100ML).

Needles for microinjection were prepared on a needle puller (Sutter Instrument P-87) using borosilicate glass capillaries with an outer diameter of 1 mm, an inner diameter of 0.58 mm and an internal filament (Warner Instruments G100F-4). Needles were bevelled at an angle of 30° with a Microelectrode beveler (Sutter Instrument BV-10). Needles were then loaded using MicroloaderTM tips (Eppendorf 5242956003), and inserted into the embryo’s anterior pole, to avoid damaging the embryo proper.

The injections were performed with a Femtojet microinjector (Eppendorf 4i) pressurised by nitrogen gas. A footswitch was used to control the duration of the injection. To ensure consistency in the volume of injection, the extent of cytoplasmic efflux from the egg was used as an indicator.

Capped mRNA synthesised from *pUC57[G43-mNeonGreen]*, *pUC57[Tc-H2Av-mScarlet-I]*, and *pCS2+[Tc-sqh-GFP]*, and dsRNA synthesised from *pGEM[Tc-twi]* and *pCR4[Tc-string]* was used for microinjection experiments. Four different mRNA and dsRNA mixes were used for injections:

- *sqh-GFP* condition: 2 µg/µL *Tc-sqh-GFP* + 1 µg/µL *Tc-H2Av-mScarlet-I* in ddH_₂_O (total concentration 3 µg/µL)
- Control condition: 2 µg/µL *G43-mNeonGreen* + 1 µg/µL *Tc-H2Av-mScarlet-I* in ddH_₂_O (total concentration 3 µg/µL)
- *Tc-twi-KD* condition: 2 µg/µL dsRNA targeting *Tc-twi* + 1 µg/µL *G43-mNeonGreen* in ddH_₂_O (total concentration 3 µg/µL)
- *Tc-string-KD* condition: 2 µg/µL dsRNA targeting *Tc-string* + 1 µg/µL *G43-mNeonGreen* in ddH_₂_O (total concentration 3 µg/µL)

Each mix was freshly prepared and centrifuged before injection to ensure homogeneity and prevent clogging of the microinjection needle. Embryos were injected with the RNA mix 2-3 hours after egg laying for all conditions, except for the *Tc-string-KD* condition in which the embryos were injected 6 hours after egg laying to avoid disrupting previous cell division rounds.

Following injections, the coverslip was turned onto the oxygen-permeable membrane of a Lumox tissue culture dish (Sarstedt #94.6077.305), as previously described^96^. The coverslip is as wide as the dish, so it can rest on the short plastic lip surrounding the membrane, to avoid squishing the embryos. Oil mix was added as needed to fill the space between the coverslip and the Lumox membrane and avoid generating excessive pressure on the embryos. Post-injection, embryos were kept at 30°C within a humid chamber, created by placing damp tissue paper into a plastic container with a lid. The Lumox plates were then brought directly to the microscope for imaging.

### *TRIBOLIUM* EMBRYO LIVE IMAGING

Live imaging was carried out at the Advanced Light Microscopy Facility (EMBL, Heidelberg). Embryos were imaged on a Leica Stellaris 8 with temperature regulated enclosure set at 32°C. Image stacks of 15 or 18 focal planes (z-step of 2 μm) were taken with a HC PL APO CS2 20x/0.75 multi-immersion objective. Embryos injected with the *Tc-sqh-GFP* and *Tc-H2Av-mScarlet-I* mRNA mix were imaged every 10 minutes. For all other conditions, embryos were imaged every 4 minutes. Acquired images were 8-bit with a physical pixel size of 0.29 μm x 0.29 μm, a 512 x 1024 scan format and a 1.5 μs/pixel dwell time. Image dimensions were 195.45 x 107.67 µm, and Z-stack step size of 1 μm. To establish collection windows, the Smart Setup tool within the manufacturer software was used. Both channels were collected simultaneously. A notch filter was used to remove the autofluorescence caused by the vitelline envelope in the green channel.

As it is not possible to establish dorsoventral orientation in blastoderm embryos, at least 10 embryos were imaged simultaneously in each session. Only time lapses in which the embryo was ventrally oriented and the midline corresponded to the middle of the image height (on average 1 in every 10), were further analysed.

### IMAGE PROCESSING AND 3D SEGMENTATION

For analysis of mesoderm cells in fixed embryos, the mesoderm region was determined based on HCR staining of *(Tc-)twi* and *(Tc-)sna*. For *Drosophila*, HCR images were analysed in Fiji^97^. First, a median filter (r = 2) was applied to the *(Tc-)twi* channel (labelling the mesoderm). Next, the image was converted to 8-bit and automatically thresholded using the Otsu’s method. The resulting mask identifies the area of the image containing mesodermal cells. This mask was then applied to identify mesoderm nuclei signal in the DAPI channel using the “AND” function in the Image Calculator menu in Fiji. For *Tribolium*, the signal was not uniform enough to enable automatic extraction of mesodermal mask. Instead, we manually selected the region based on the mesodermal and anteroposterior genes HCR signal.

The mesodermal nuclei were then segmented in 3D using the machine learning framework Cellpose 2.0^98^, run on Python 3.8.12^99^. For each image, the software was automatically calibrated to the nuclear size, and nuclei were segmented using the cytoplasm 2.0 model, which worked better than nuclear models for our dataset. Other parameters were kept to their default values. Wrongly segmented masks were manually deleted within the Cellpose GUI. Mesodermal nuclei masks were then loaded into Fiji to extract centroid coordinates for each mask. These were computed using the 3D Centroid feature in the MorphoLibJ plug-in^100^.

To segment mesodermal cells in 3D, we used the LimeSeg plug-in in Fiji^97^. Nuclear centroids were used as seeds for the segmentation of cell outlines, marked by the phalloidin signal. To do so, we used a script kindly provided by Erik Clark (University of Cambridge), which is a modified version of the script DemoVescicleSegmentation.groovy (https://github.com/NicoKiaru/LimeSeg/blob/master/src/main/resources/script-templates/LimeSeg/DemoVesicleSegmentation.groovy). The script reads a text file containing nuclear centroid coordinates, and it adds a spherical ROI centred around the nuclear centroid to the phalloidin channel. It then launches the Limeseg spherical segmentation function. The algorithm allows the spherical seeds to grow simultaneously following dynamics inspired by lipid membranes behaviours, until they reach local intensity peaks. Parameters were set as follows: d_0 = 3.5, f_pressure = 0.015, z_scale = 3.376, range_in_d0_units = 1.5. The optimisation process was halted after 10 minutes once the masks achieved sufficient convergence. For verifying segmentation accuracy, a custom MATLAB (versions R2022bR2024a)^101^ script by Dr. Wolfram Pönisch was employed to transform the triangulated meshes into masks. The 3D cell masks channel was fused with the corresponding phalloidin channels in Fiji. Incorrect masks (which mainly comprised of masks which outgrew the embryo surface and filled the background of the image) were eliminated using the Label Edition plug-in’s GUI in MorphoLibJ. This procedure resulted in directories containing .ply files of accurately segmented cell masks.

### 3D CELL SHAPE ANALYSIS VIA SPHERICAL HARMONICS AND DIMENSIONALITY REDUCTION ALGORITHMS

The point clouds contained in the .ply files were then subjected to spherical harmonics decomposition and dimensional reduction using a previously described framework^32^. In brief, the point clouds obtained from cell segmentation were triangulated (using the MATLAB function MyRobustCrust.m, see https://github.com/markeroon/matlab-computer-vision-routines/blob/master/third_party/MyCrust100809/MyRobustCrust.m) so that a 3D shape is represented as a set of surface points (called vertices) linked by edges and forming triangles (called faces). Next, surfaces with holes representing incomplete segmentation were removed. Cells were then aligned along their longest axis as follows. For alignment, we calculated the (volumetric) covariance matrix of the centred cell volume and defined the new axes from its eigenvectors. The initial direction of the cell alignment was chosen such that the third moment ⟨*x*^3^⟩ along the new *x*-axis was positive. The direction of the medium axis was chosen such that ⟨*x*^2^*y*⟩ > 0. After alignment, cell shapes were simplified using the MATLAB function reducepatch() to reduce the number of faces per cell to 2000. This led to a significant acceleration of the following spherical harmonics decomposition. The rotated *x*, *y*, and *z* components of the vertices were extracted and mapped onto a sphere using an area-preserving flattening of the surface points of the cell^102^. This method allows for the description of shapes that are not star-convex, as it extracts three distributions on the surface of a sphere, one for each dimension *x*, *y*, *z*. For each cell, spherical harmonics decomposition were performed

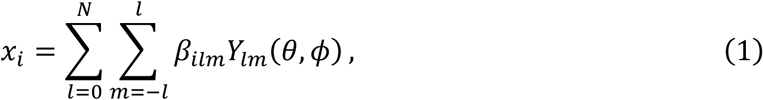

with *i* = 1,2,3 and where the coordinates *x*_1_, *x*_2_, *x*_3_ correspond to *x*, *y*, *z*. *Y_lm_*(*θ, φ*) are the spherical harmonics functions with the angles *θ* and *φ*. The coefficients *β_ilm_* are the spherical harmonics amplitudes which are unique shape descriptors. We also introduced *N*, the highest degree of considered spherical harmonics functions. We found that the contribution of spherical harmonics functions above 20 do not significantly contribute to the total cumulative variance of spherical harmonics coefficients (see Fig. S1B)

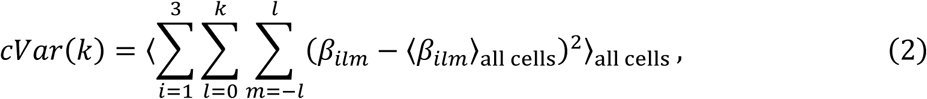

and hence do not contribute to a PCA as well. In our case we used *N* = 40 orders, totalling 5043 coefficients (1681 in each of the three dimensions *x*, *y*, *z*). After decomposing all cell shapes, we estimated their errors by computing the distances of the vertices of the original shape and the surface reconstructed from its spherical harmonics coefficients and viceversa using the point2trimesh() function in MATLAB (https://www.mathworks.com/matlabcentral/fileexchange/52882-point2trimesh-distance-between-point-and-triangulated-surface). We then calculated the mean and maximum error for each cell shape and only considered shapes with an average error below 0.1 µm and a maximum error below 2 µm.

For morphospace representation, the spherical harmonics datasets were then subjected to dimensionality reduction using either PCA or t-Distributed Stochastic Neighbor Embedding (tSNE). For the PCA, we calculated the number of principal components (PCs) *N_PC_* to account for 90% of the total variance (Fig. S1C). Mean cell shapes (see Fig. 1 and 2) and background shapes in Fig. 1, S1, 2, S2 and S3 were reconstructed based on principal component values. To define distances in the morphospace in a meaningful way, we computed the Euclidean distance in the morphospace for two cell shapes *m* and *n*

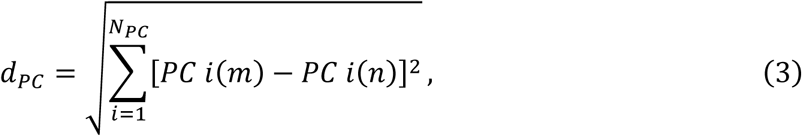

and compared it to the mean error between the two shapes.

To generate bagplots^103^ (see Fig. S1, 2, S2 and S3), we employed the MATLAB LIBRA toolbox^104^. tSNE was utilised with a perplexity of 30.

Three datasets were used to build the morphospaces:

1. The global morphospace included all cells from 12 embryos in *Drosophila* and 14 embryos in *Tribolium* (Fig. 1 and S1), with an average number of cells per stage of 316 +/- 170 for *Drosophila*, and 291 +/- 75 for *Tribolium*;
2. A normalised morphospace included all cells from all the embryos in *Drosophila* and *Tribolium* normalised so that each cell had the same volume (Fig. 2 and S2);
3. A focused morphospace including 6 embryos for each species, two for each stage (blastoderm, ventral furrow, tube) (Fig. 2 and S3).

It is important to note that in the *Drosophila* embryos 3 and 4 (ventral furrow stage) only the mesodermal cells that apically constrict where included in the analysis, as the most dorsal mesodermal cells display a different behaviour by expanding their apex^105^.

### CELL DIVISION ORIENTATION

For live imaging of cell division in *Tribolium*, 6 time lapses (three ectoderm views and three mesoderm views) were selected from four independent experiments in embryos injected with *G43-mNeonGreen* RNA (labelling cellular membranes) and *H2Av-mScarlet-I* RNA (to mark chromatin), over 11 time steps (44 minutes total). Each time lapse was manually cropped so that Frame = 1 (Time = 0 minutes) corresponded to the completion of cellularisation, which can be clearly identified as the cellular membranes condense as bright puncta at the basal side of each cell^37^. We then used Fiji to extract the H2Av channel, apply a median filter (radius = 5 µm), and enhance contrast. Nuclei and mitotic chromosome masses were then segmented using the Object Classification workflow in Ilastik^106,107^, and exported as Object Identities .tiff files, in which each nucleus in each frame is labelled as a mask with a unique pixel value as its identifier. The masks were then merged with the original H2A signal in Fiji to form an RGB stack, to check for accuracy. For each cell division, the two daughter cells were manually selected using the multipoint tool in the first frame in which they appeared as two separate objects. The pixel values of the corresponding two chromosome masks were saved in a .csv spreadsheet as 3D coordinates, together with the time frame, the sample name and the integer value of each mask.

All subsequent analyses described in this article were performed in Python (v3.8.18)^99^ using the NumPy^108^, pandas^109^, SciPy^110^ and scikit-image^111^ libraries within Jupyter Notebooks. Data visualisation employed Matplotlib^112^ and Seaborn^113^. Specific packages are cited throughout the text if not included in the above list.

For determining division orientation, we used a Jupyter Notebook to extract chromosome mass centroids from the Object Identities .tiff files using the unique pixel value for each post-division pair listed in the accompanying .csv tables. For each nucleus at each time point, voxel coordinates corresponding to its label (pixel value) were identified, and the centroid was calculated as the mean of voxel indices scaled by its physical dimensions (0.284 × 0.284 × 2 µm per pixel). Pairwise displacement vectors between associated chromosome masses were then computed, and the polar angle θ (relative to the z-axis, with reversed convention) was derived using trigonometric relationships. For visualisation, θ values from mesoderm and ectoderm were plotted as scatter points over time, with mean trajectories and mean ± SEM per time bin (smoothed by a Gaussian filter, σ = 2) overlaid to highlight temporal trends.

### LIVE IMAGE PROCESSING, TRACKING, AND FEATURE EXTRACTION

For live imaging of *Tribolium* mesoderm during gastrulation, three time-lapses were selected per condition. Embryos were injected as described above with mRNA encoding either *G43-mNeonGreen* and *H2Av-mScarlet-I* (control), or *G43-mNeonGreen* together with dsRNA targeting *Tc-twi* or *Tc-string*. Time-lapse images were cropped and aligned so that the first frame corresponded to the completion of cellularisation, and that the left side of the image corresponded to the region posterior to head formation.

If the nuclear channel was present (only in control embryos), the channel was processed in Fiji using a median filter (r = 2) before generating projections. For the membrane channel, an average projection was generated using the LocalZProjector Fiji plug-in^114^ with the following parameters: binning = 4, method = max of std, neighbourhood size = 16, gaussian pre = 0.70, median post = 30, method = MIP. This plug-in produced two outputs: a projected 2D stack and a height map indicating the height of pixels in the original z-dimension. The height map was later used for morphological corrections.

Automatic segmentation of the apical membranes was performed on the membrane 2D projection using the Tissue Analyzer plug-in^115^ with a watershed method (strong blur = 3.5, weak blur = 1.0). The segmentation was manually verified for accuracy. The resulting binary masks were imported into Ilastik for manual tracking. This tracking process produced:

- A .tiff file, where each tracked object was assigned a unique pixel value that remained consistent across frames;
- A .csv file, containing cell division data linking mother cells to daughter cells and specifying the frame in which each division occurred.

For out-of-plane divisions and extrusions, division and extrusion times were annotated manually in a .txt file. The original z-stack was inspected to confirm these events. Manual tracking overall ensured that only correctly segmented mesodermal cells were included in the following analysis.

All .tiff and .txt files were processed in a Jupyter Notebook. Binary masks were converted into labelled objects using ndimage.label (SciPy), with pixel values corresponding to tracking IDs.

For control embryos, a set of morphometric features were extracted from the data. A detailed explanation of feature calculation, scale and corrections can be found in Table S1. The following geometrical features were extracted for each object using the regionprops function (scikit-image) and normalised to microns using the pixel size of the image (0.284 µm^2^/pixel): area, convex area, major axis length, minor axis length, centroid x, centroid y, and orientation. To correct for height distortions, features were adjusted using the height map from LocalZProjector by adapting the original *DeProj* code from MATLAB to Python. Additional features (eccentricity, extent and solidity) were computed from corrected values (see Table S1).

### LINEAGE ASSIGNMENT, DENSITY MEASUREMENTS, AND EXTRUSION TRACKING

The tracking data was processed to assign division times to each object and determine lineage identity. A new column, normalised time, was created by subtracting the division time (from the .csv file output of the manual tracking workflow in Ilastik, or manually assigned for out of plane divisions by inspecting the z-stacks) from the absolute time. This time was therefore normalised to the time-point of division. This transformation ensured that normalised time = 0 corresponded to the last time point when the cell mother was a unique object. Each ventral blastoderm cell was classified as either a mother (pre-division) or a daughter (post-division) cell. In total, 93 mother and 165 daughter cells were analysed in Sample 1, 99 mothers and 176 daughters in Sample 2, and 58 mothers and 92 daughters in Sample 3. From our manual tracking observation, ventral blastoderm cells divide only once between cellularisation and gastrulation.

Three density parameters were calculated: apical density, relative area and nuclear crowding (see details in Table S1). To ensure accurate measurements at the mesoderm boundary, each tracked cell at each time point was matched to segmentation outputs generated with TissueAnalyzer (apical cell masks) and Cellpose (nuclear masks). The tracking dataset contained only the cells that were manually followed over time, whereas the segmentation outputs contained masks for all cells present in the field of view at each time point. Matching the tracked cells to these full segmentation datasets therefore allowed density measurements to be calculated relative to the complete local cell population rather than only the subset of tracked cells. This ensured that measurements accounted for all neighbouring cells within the analysis window, provided the window did not extend beyond the image boundaries. Apical density and relative area were measured in a window of radius = 20 µm centred around the centroid of the cell apex mask, while nuclear crowding was measured in a window of radius = 10 µm centred around the nucleus centroid. These measures were defined after testing windows of various radii (from 5 to 25 µm) to roughly include the closest neighbours at each time point, adjusted for each category. As the cell area changes dynamically, and the cell neighbours are continuously exchanging due to the asynchronous nature of divisions in the *Tribolium* mesoderm, it was not possible to automatically track the cell neighbours at each time point, and we used measurements within a fixed window as an approximation instead.

For analysis of feature dynamics with respect to cell extrusion, extrusion time was manually assigned from the .csv file recorded during tracking, and defined as the last time point at which an extruding cell was observed in the apical projection.

Cells were classified as anterior or posterior based on their average centroid x position, using half the sample length as the boundary between the two regions. This does not necessarily match “anterior” or “posterior” embryonic regions in a canonical sense, but rather the relative position of the mesodermal cells found in the field of view that was imaged and analysed.

Processed data were saved as .csv files for further analysis. Visualisations, including time series plots and density comparisons, were generated using Seaborn and Matplotlib. Statistical comparisons between independent groups were performed using the non-parametric Mann–Whitney U test (two-sided), implemented in SciPy.

### FURROW DEPTH MEASUREMENTS

Furrow depth measurements were performed manually for three control embryos. Each image stack was cropped to include the mesodermal region in antero-posterior and dorso-ventral direction, and aligned so that time=0 corresponded to the completion of cellularisation. The stack was divided into three equally spaced regions along the anteroposterior axis (left-right axis of the image), re-sliced, and measured using the line tool in Fiji. The depth of the furrow was manually determined at each relevant position within the re-sliced stack. Before furrowing began, the depth was recorded as 0.

### RANDOM FOREST CLASSIFIER AND SHAP VALUES ANALYSIS OF PREDICTIVE FEATURES

To identify features predictive of division orientation, a Random Forest Classifier (scikit-learn) was trained with the geometrical and tissue density features described above (see also Table S1), with Shapley Additive Explanations (SHAP) values used to assess specific feature importance. SHAP values were computed on Random Forest Classifiers using the Shap package^116^, which provides model-agnostic explanations for predictions. SHAP summary and dependency plots were generated in the Shap package. The feature set for the SHAP analysis included: area, apical density, relative area, eccentricity, extent, orientation, solidity, and nuclear crowding.

For the random forest classifier analysis, we generated a dataset containing all mother cells’ features over 28 minutes preceding each division, with the features averaged in bins over two time points as explained in the main text. This dataset was then split into a training set (60%), validation set (20%), and test set (20%). We used the RandomForestClassifier package from Scikit-learn. For each time bin, a separate model was trained and evaluated using F1-scores, which report the harmonic mean of precision (the proportion of correctly predicted positive cases among all predicted positives) and recall (the proportion of correctly predicted positive cases among all actual positives).

To further explore the relationship between feature importance and developmental time, SHAP values were computed for a random forest model trained specifically on the time bin most predictive of division orientation, -12 to -8 min. SHAP values were used to determine the relative contributions of specific features to cell division orientation at this stage.

## SUPPLEMENTARY INFORMATIONS

**Figure S1.**
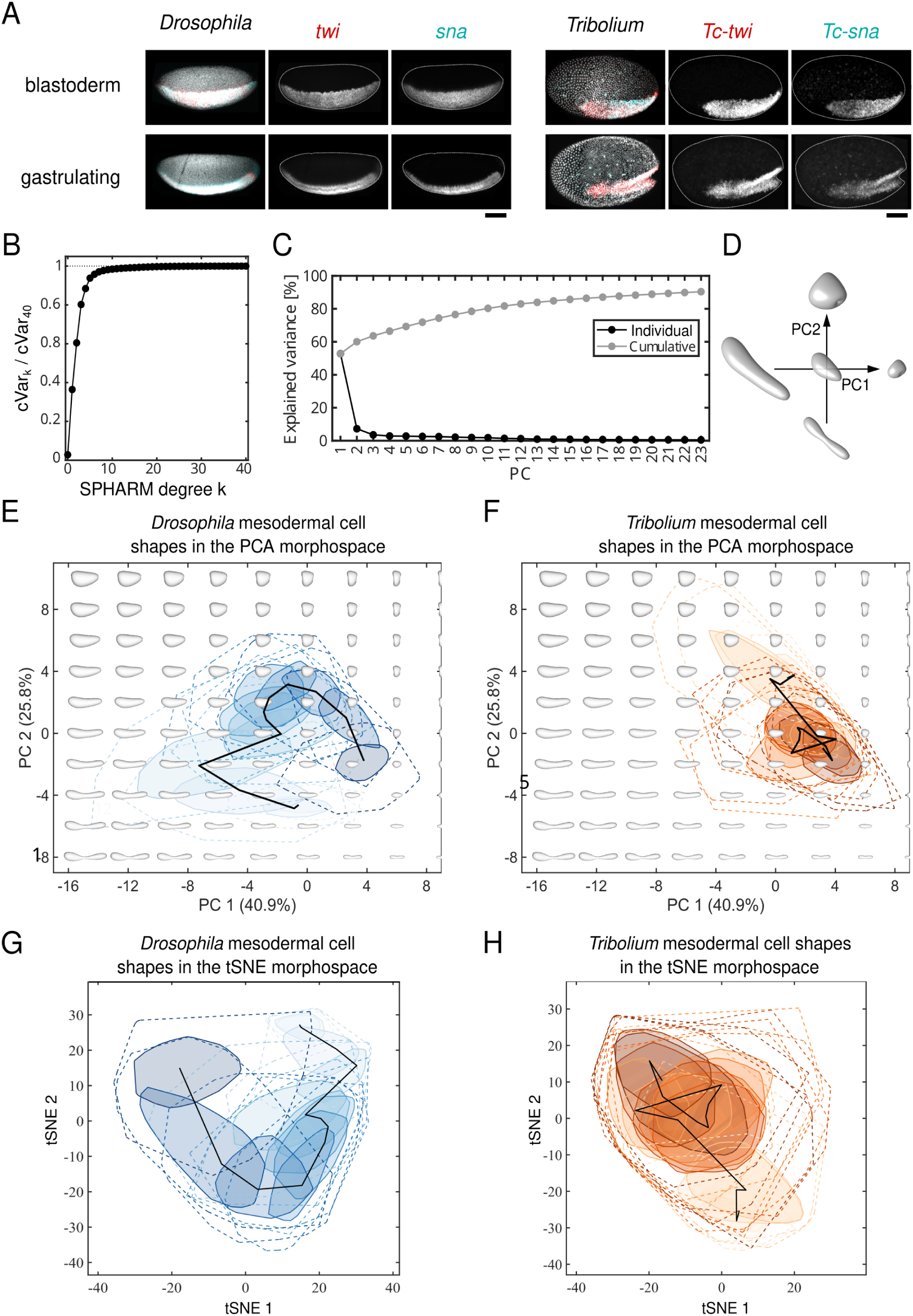
Supplementary data on cell shape trajectories during mesoderm gastrulation in *Drosophila* and *Tribolium*, related to Figure 1. (A) Staining of *twi* and *sna* in *Drosophila* embryos and their *Tribolium* homologues (*Tc-twi* and *Tc-sna*) in *Tribolium* embryos during the blastoderm (top panels) and ventral furrow (bottom panels) stages of gastrulation. Embryos were stained with DAPI to visualise nuclei and probed with HCR v3 to detect transcripts. Left panels: merged view of DAPI and HCR signal. Scale bars, 100 µm. (B) Total cumulative variance of spherical harmonics coefficients (Eq. 1). (C) Individual and cumulative percentage of cell shape variance in the morphospace constructed with all the cells segmented in *Drosophila* and *Tribolium*, explained by the first 23 PCs, corresponding to 90% of the total variance. (D) Reconstructed cell shape changes reflected by changes in PC1 and PC2. The central shape corresponds to PC1 = PC2 = 0. (E and F) Bag-plot representation of mesoderm 3D cell shapes from *Drosophila* (E) and *Tribolium* (F) embryos undergoing gastrulation, plotted in the PC1-PC2 morphospace. Polygons represent bagplots: the inner polygon (bag) contains 50% of points, the outer polygon (fence) corresponds to the convex hull of all data. Black line: cell shape trajectory as defined in Fig. 1E,F. (G and H) t-SNE embedding of *Drosophila* (G) and *Tribolium* (H) mesoderm cell shapes. Bag-plots are shown as in E,F. For (E-H), each bag-plot corresponds to one embryo, and darker shades correspond to later developmental times, as in Fig. 1E and F.

**Figure S2.**
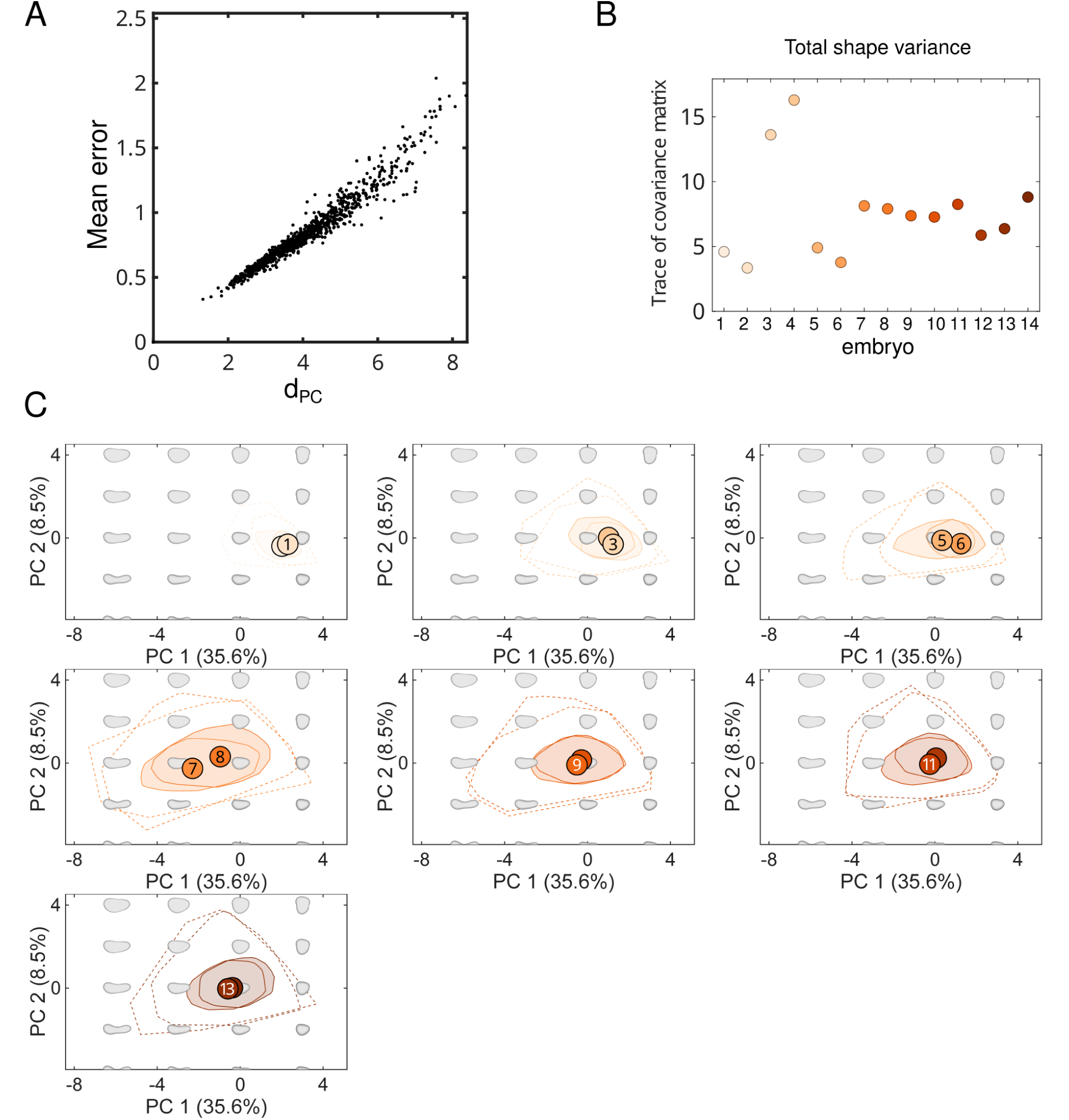
Analysis of *Tribolium* mesoderm cell shapes normalised for volume shows that cell shape variability peaks at ventral furrow formation stage, related to Figure 2. (A) Comparison of distances in morphospace (see Eq. 3) and mean surface distance between cell shapes (see Methods). Here, we chose 1000 pairs of cells used to generate the distances (see Methods). (B) Total variance of all spherical harmonics functions for each stage after cell volume normalisation (see Eq. 2 and Methods). (C) Bag plot representation of mesoderm 3D cell shapes from *Tribolium* embryos undergoing gastrulation, plotted in the PC1-PC2 morphospace. Cell volume was normalised so that all cells have the same volume. Each graph has plotted data from two embryos at the same developmental stage, with darker shades corresponding to later developmental times, as in Fig. 1F. Bag plot representation: the inner polygon (bag) contains ≤50% of points, the outer polygon (fence) corresponds to the convex hull of all data.

**Figure S3.**
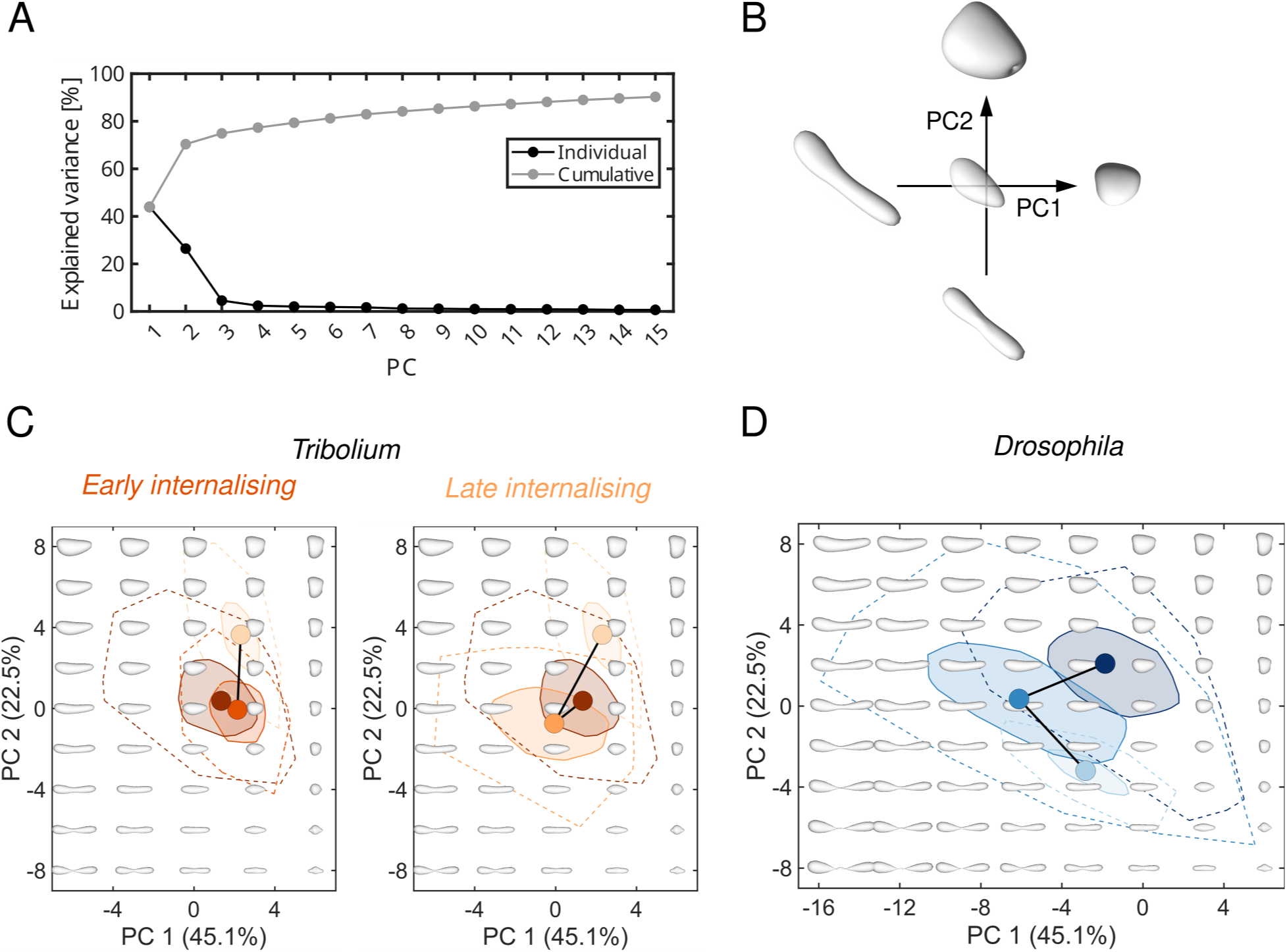
Supplementary data for the focused morphospace analysis of mesodermal cell shapes in *Tribolium* and *Drosophila* during ventral furrow formation stage, related to Figure 2. (A) Individual and cumulative percentage of cell shape variance in the minimal dataset of *Drosophila* and *Tribolium* embryos at blastoderm, ventral furrow and tube stages (as explained in Fig. 2) explained by the first 15 PCs (corresponding to 90% of the total variance). PC1 and PC2 were used to construct the focused morphospace. (B) Reconstructed cell shape changes reflected by changes in PC1 and PC2 in the focused morphospace. The central shape corresponds to PC1 = PC2 = 0. (C and D) Bag-plot representation of mesoderm 3D cell shapes from *Tribolium* (C) and *Drosophila* (D) embryos during ventral furrow formation, plotted in the PC1-PC2 focused morphospace from Fig. 2. Polygons represent bagplots: the inner polygon (bag) contains ≤50% of points, the outer polygon (fence) corresponds to the convex hull of all data. Black line: cell shape trajectory.

**Figure S4.**
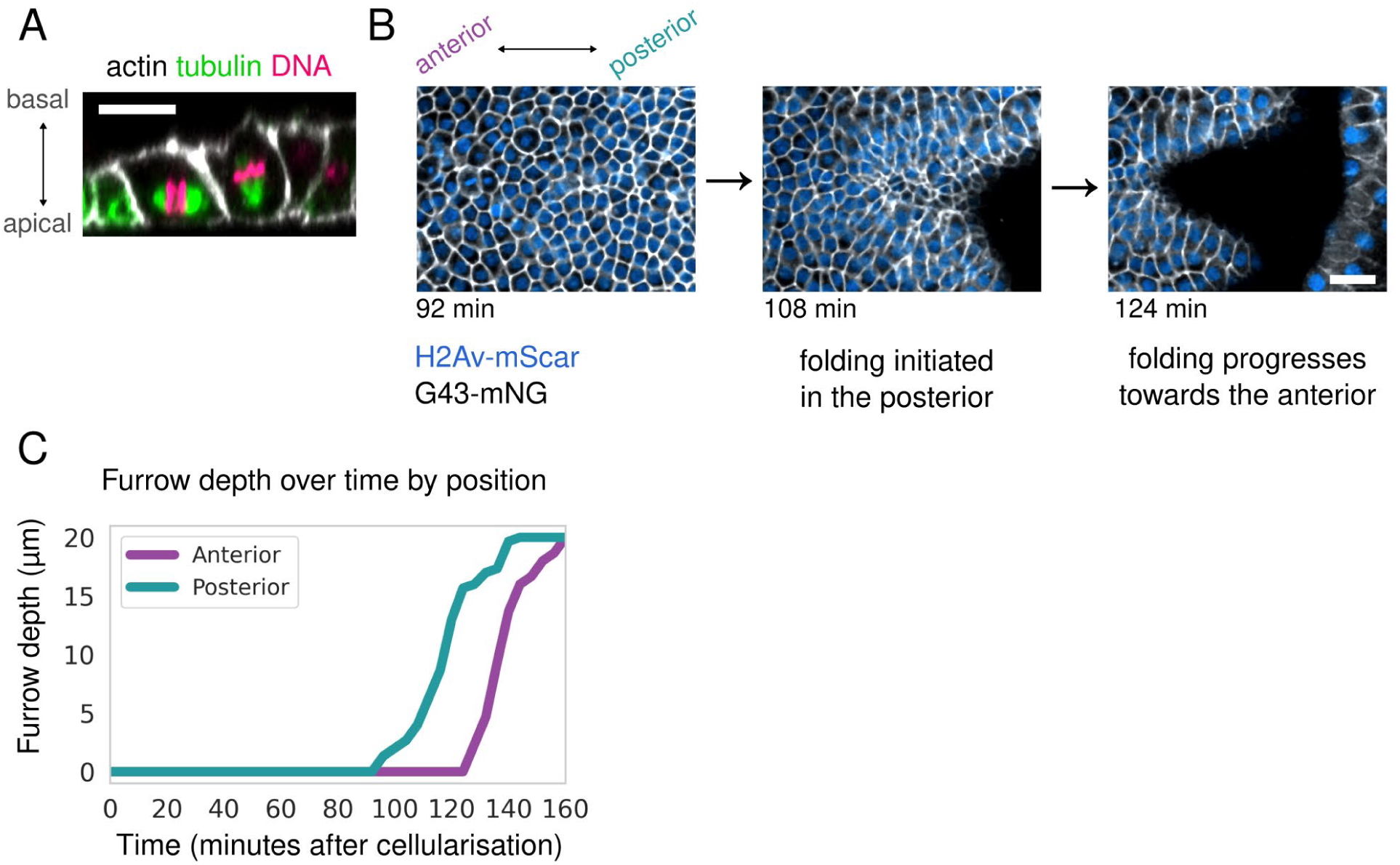
Additional information for the analysis of the location and timing of out-of-plane divisions in the *Tribolium* mesoderm, related to Figure 3. (A) Two dividing cells in the *Tribolium* mesoderm displaying spindles positioned in the plane (cell on the left) and out of the plane (cell on the right) of the epithelium. The embryo was stained for acetylated alpha tubulin to visualise microtubules (green), phalloidin to visualise actin (greyscale) and DAPI to visualise DNA (magenta). The image is a cross-section of a confocal stack, averaged across 4µm. Scalebar, 10µm. (B) Apical view (average projection of grazing section of ∼4µm) of a confocal microscopy timelapse of the *Tribolium* mesoderm transiently expressing H2Av-mScarlet to mark DNA (blue) and G43-mNG to label the plasma membrane (greyscale) showing that ventral furrow formation and mesoderm invagination proceed from the posterior towards the anterior side of the embryo. Time is given in minutes post cellularisation. Scalebar, 20µm (C) Furrow depth over time averaged over 3 embryos, plotted by position (anterior, middle, posterior, defined at three evenly spaced locations along the antero-posterior axis). Furrow depth was measured manually in cross-sections for each time-lapse.

**Figure S5.**
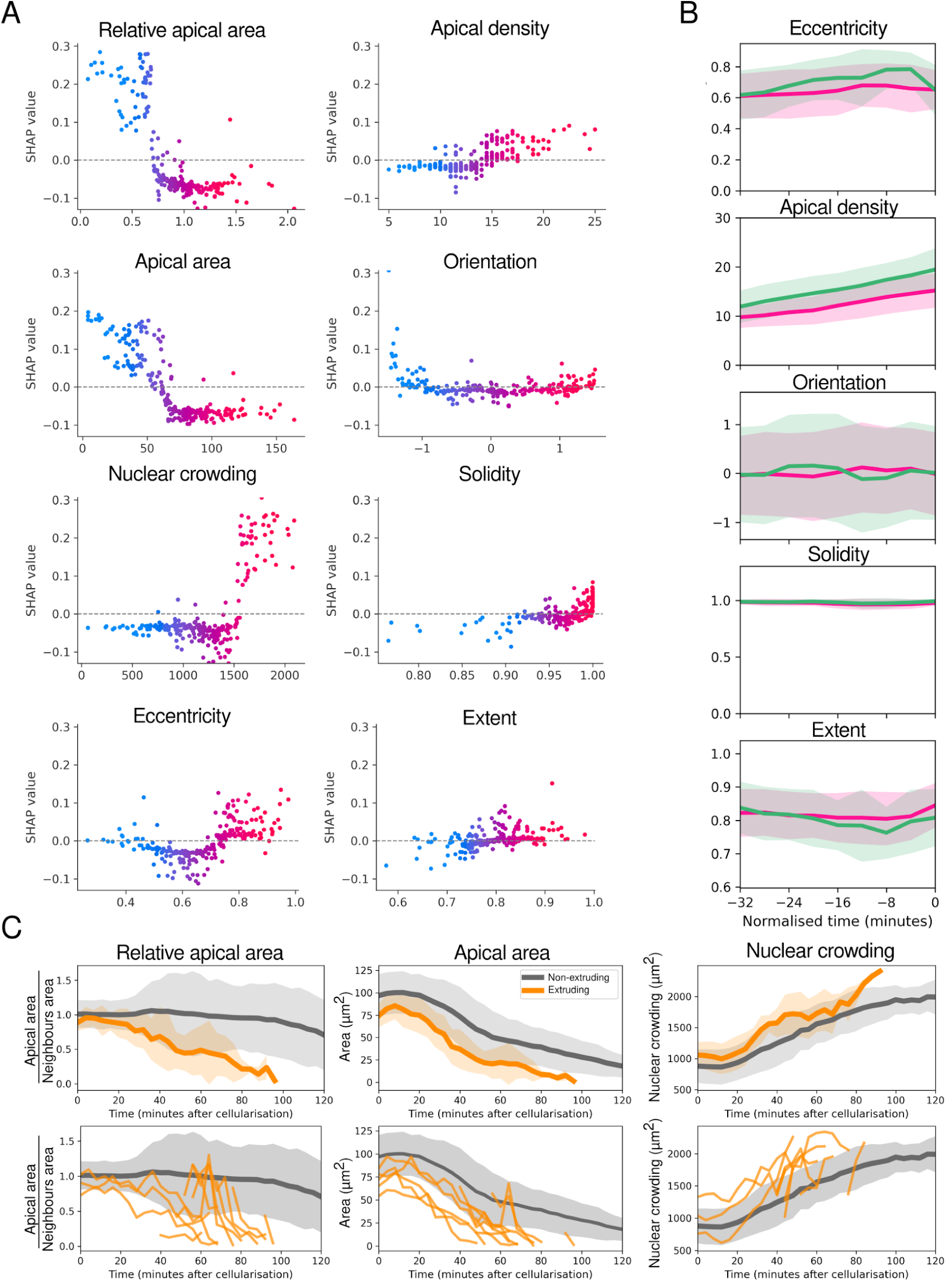
Additional data for the interpretable machine learning analysis of local cellular features predicting division orientation, and dynamics of key features for cells undergoing extrusion, related to Figure 4. (A) SHAP dependence plots for the features used to predict division orientation using the Random Forest Classifier, showing the contribution of each feature to model predictions. Refer to Table S1 for the unit used for each feature on the x axis. Each point represents one cell, color-coded by the feature value (gradient from blue = low values to red = high values). Higher SHAP values indicate a stronger contribution of that feature toward an out-of-plane division prediction, whereas lower (more negative) SHAP values indicate a contribution toward an in-plane division prediction. (B) Dynamics of the 5 less predictive local cellular features (eccentricity, apical density, orientation, solidity, extent) over normalised time for in plane (pink, *n* = 189) and out-of-plane (green, *n* = 61) divisions. *N* = 3 embryos were analysed. Solid line: mean, shaded area highlights standard deviation. Refer to Table S1 for the unit used for each feature on the y axis (C) Dynamics of apical area, relative apical area, and nuclear crowding in extruding cells compared to all mesoderm cells. Top row: mean ± standard deviation; bottom row: individual cell tracks of 10 randomly selected cells. *n* = 628 cells across 3 embryos, with a total of 21 extruding cells (3% of all cells extrude).

**Figure S6.**
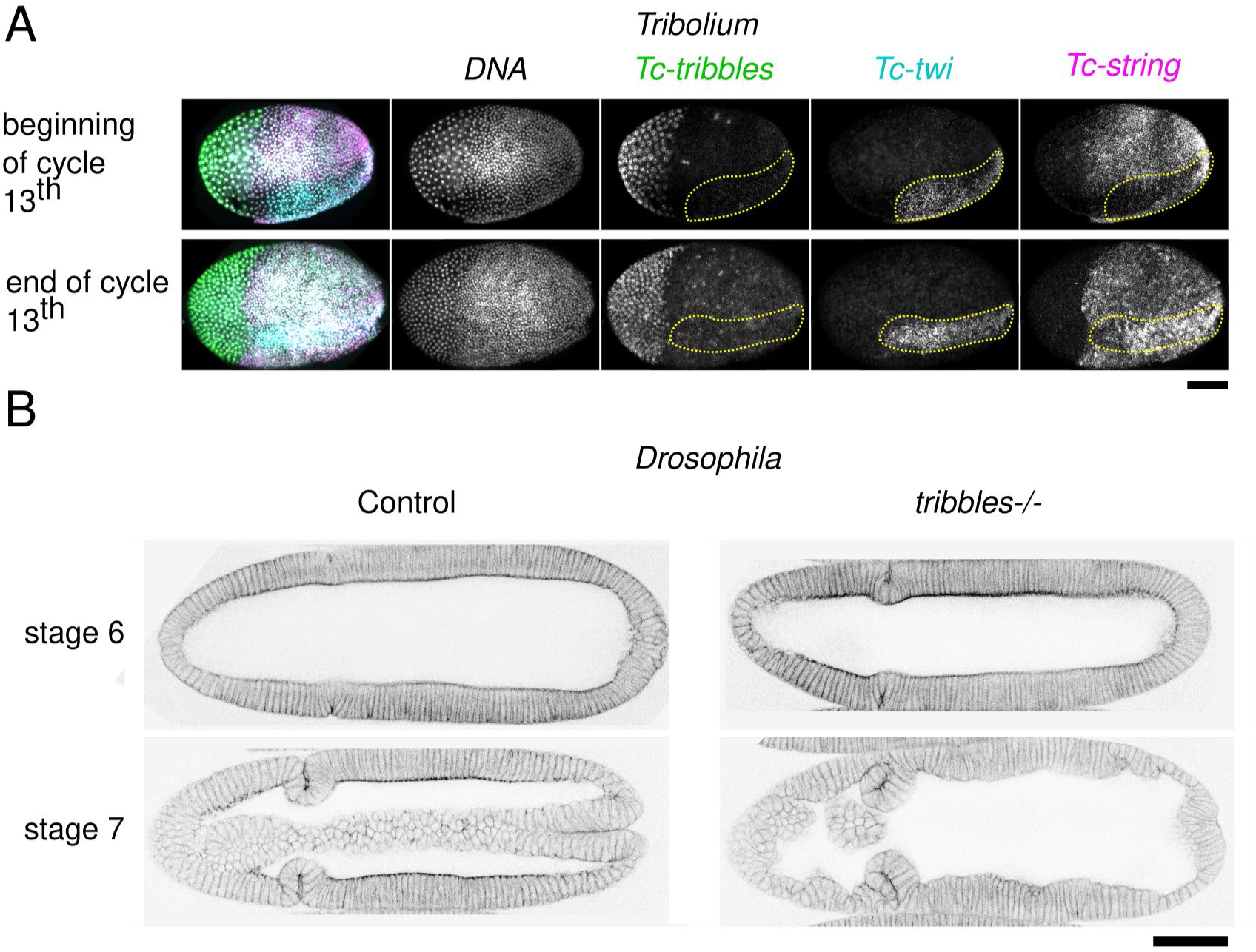
*Tribolium Tc-tribbles* and *Tc-strin*g expression in the mesoderm, and *Drosophila tribbles* mutant embryo staging, related to Figure 5. (A) Expression of *Tc-tribbles*, *Tc-twi* and *Tc-string* during cycle 13. The beginning and end of cycle 13 are shown for each condition. *Tc-tribbles* expression is restricted to the extraembryonic tissue. *Tc-string* is first expressed in the ectoderm, then in the ectoderm and mesoderm. Yellow dashed lines mark the mesodermal region. Scale bar, 100 µm. (B) Representative images of control and *tribbles Drosophila* embryos at two successive stages of mesoderm invagination, corresponding to embryos from Fig. 5. Embryos were stained for phalloidin (inverted contrast is displayed). Scale bar, 40 µm.

**Table S1.**
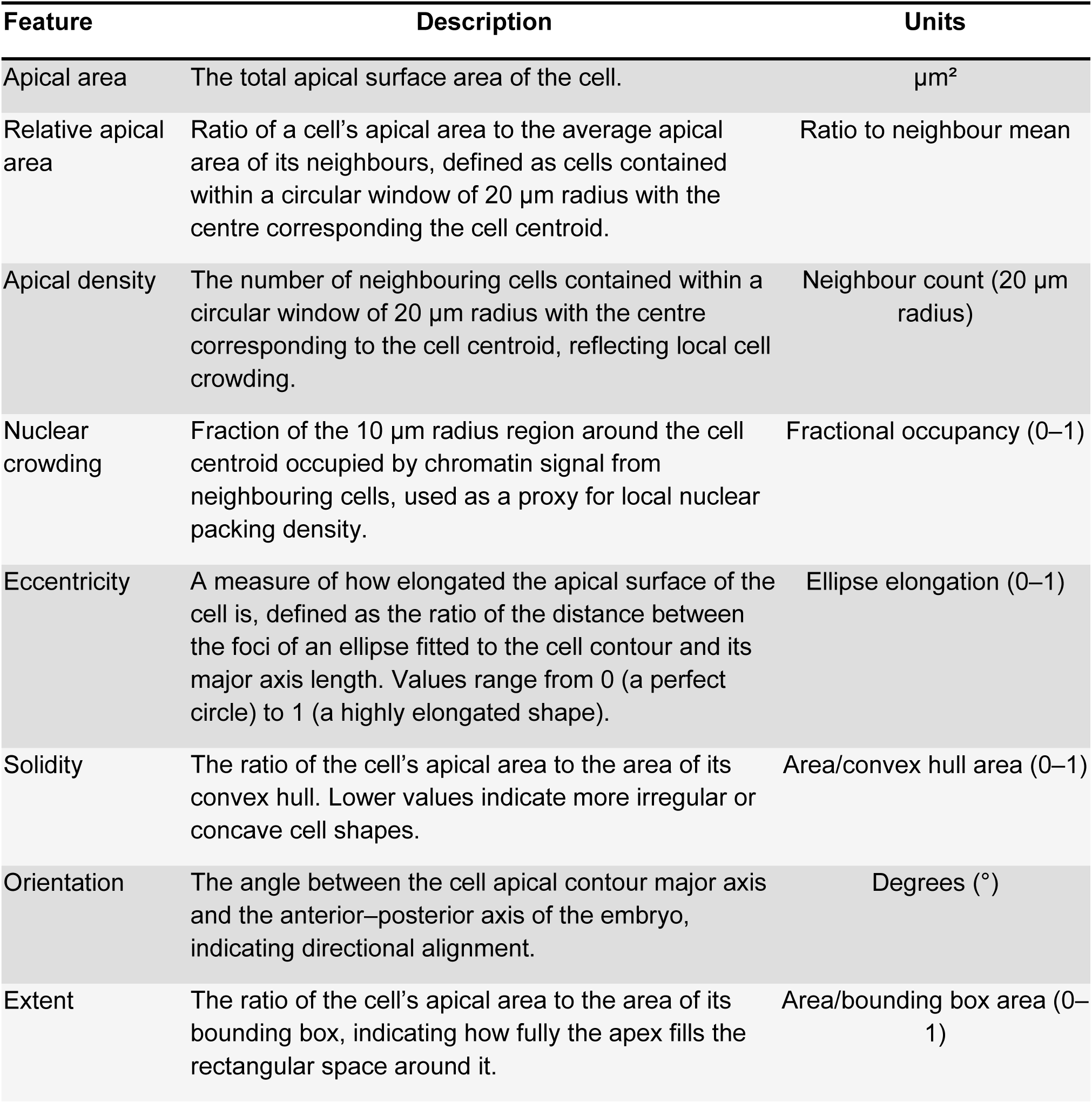
Morphodynamic features.

| Feature | Description | Units |
| --- | --- | --- |
| Apical area | The total apical surface area of the cell. | $\mu\text{m}^2$ |
| Relative apical area | Ratio of a cell's apical area to the average apical area of its neighbours, defined as cells contained within a circular window of 20 $\mu\text{m}$ radius with the centre corresponding the cell centroid. | Ratio to neighbour mean |
| Apical density | The number of neighbouring cells contained within a circular window of 20 $\mu\text{m}$ radius with the centre corresponding to the cell centroid, reflecting local cell crowding. | Neighbour count (20 $\mu\text{m}$ radius) |
| Nuclear crowding | Fraction of the 10 $\mu\text{m}$ radius region around the cell centroid occupied by chromatin signal from neighbouring cells, used as a proxy for local nuclear packing density. | Fractional occupancy (0–1) |
| Eccentricity | A measure of how elongated the apical surface of the cell is, defined as the ratio of the distance between the foci of an ellipse fitted to the cell contour and its major axis length. Values range from 0 (a perfect circle) to 1 (a highly elongated shape). | Ellipse elongation (0–1) |
| Solidity | The ratio of the cell's apical area to the area of its convex hull. Lower values indicate more irregular or concave cell shapes. | Area/convex hull area (0–1) |
| Orientation | The angle between the cell apical contour major axis and the anterior–posterior axis of the embryo, indicating directional alignment. | Degrees ($^\circ$) |
| Extent | The ratio of the cell's apical area to the area of its bounding box, indicating how fully the apex fills the rectangular space around it. | Area/bounding box area (0–1) |

## SUPPLEMENTARY VIDEOS TITLES AND CAPTIONS

**Video S1 Myosin signal in *Tribolium* mesoderm, related to Figure 2**. Time-lapse of a *Tribolium* embryo injected with *squash-GFP* RNA to visualise non-muscle myosin II. The mesoderm apical domain shows a burst of signal before undergoing folding. Average projection of grazing section of 4∼µm. Scale bar, 20 μm.

**Video S2. Out-of-plane cell division in *Tribolium* mesoderm, related to Figure 3**. Time-lapse of *Tribolium* mesodermal cells undergoing out-of-plane cell division. Embryos were injected with *G43-mNeonGreen* RNA to label cell membranes (grayscale) and *H2Av-mScarlet* RNA to label nuclei (cyan). Left, membranes and nuclei; right, nuclei only. Apical views are shown at the top and corresponding cross-sections at the bottom. A yellow dot tracks a mesodermal cell until division, after which the internalized daughter cell is shown in light orange and the apical daughter cell in dark orange. Average projection of grazing section of 4∼µm. Scale bar, 10 μm.

**Video S3. Mesoderm cell behaviours in control and *Tc-twi* RNAi embryos, related to Figure 5**. Time-lapse of the ventral region of *Tribolium* embryos injected with *G43-mNeonGreen* RNA to label cell membranes (grayscale). The embryo on the right was co-injected with dsRNA targeting *Tc-twi*; the embryo on the left is a control embryo. Bottom panels show the corresponding segmented and tracked cells, colour-coded according to behaviour: in-plane divisions (pink), out-of-plane divisions (green), and cell extrusion events (yellow). Average projection of grazing section of 4∼µm. Scale bar, 20 μm.

**Video S4. Mesoderm morphogenesis in control and *Tc-string* RNAi embryos, related to Figure 5**. Time-lapse of the ventral region of *Tribolium* embryos injected with G43-mNeonGreen RNA to label cell membranes (grayscale). The embryo on the right was co-injected with dsRNA targeting *Tc-string*; the embryo on the left is a control embryo. Average projection of grazing section of 4∼µm. Scale bar, 20 μm.

## Notes

### Competing Interest Statement

The authors have declared no competing interest.

### Summary of Updates

Reference fix in the method section.

## REFERENCES

1. Trinkaus, J.P. (1984). Cells Into Organs. The Forces That Shape the Embryo. Second Edition (Prentice-Hall, Englewood Cliffs).

2. Burda, I., Martin, A.C., Roeder, A.H.K., and Collins, M.A. (2023). The dynamics and biophysics of shape formation: Common themes in plant and animal morphogenesis. Developmental Cell 58, 2850–2866. 10.1016/j.devcel.2023.11.003.

3. Stern, C.D. (2004). Gastrulation: From Cells to Embryo 2004th ed. (Cold Spring Harbor Laboratory Press).

4. Nájera, G.S., and Weijer, C.J. (2023). The evolution of gastrulation morphologies. Development 150, dev200885. 10.1242/dev.200885.

5. Nakaya, Y., and Sheng, G. (2008). Epithelial to mesenchymal transition during gastrulation: An embryological view. Development, Growth & Differentiation 50, 755–766. 10.1111/j.1440-169X.2008.01070.x.

6. Solnica-Krezel, L., and Sepich, D.S. (2012). Gastrulation: Making and Shaping Germ Layers. Annual Review of Cell and Developmental Biology 28, 687–717. 10.1146/annurev-cellbio-092910-154043.

7. Francou, A., Anderson, K.V., and Hadjantonakis, A.-K. (2023). A ratchet-like apical constriction drives cell ingression during the mouse gastrulation EMT. eLife 12, e84019. 10.7554/eLife.84019.

8. Rozbicki, E., Chuai, M., Karjalainen, A.I., Song, F., Sang, H.M., Martin, R., Knölker, H.-J., MacDonald, M.P., and Weijer, C.J. (2015). Myosin-II-mediated cell shape changes and cell intercalation contribute to primitive streak formation. Nat Cell Biol 17, 397–408. 10.1038/ncb3138.

9. Williams, M., Burdsal, C., Periasamy, A., Lewandoski, M., and Sutherland, A. (2012). Mouse primitive streak forms in situ by initiation of epithelial to mesenchymal transition without migration of a cell population. Developmental Dynamics 241, 270–283. 10.1002/dvdy.23711.

10. van der Sande, M., Kraus, Y., Houliston, E., and Kaandorp, J. (2020). A cell-based boundary model of gastrulation by unipolar ingression in the hydrozoan cnidarian *Clytia hemisphaerica*. Developmental Biology 460, 176–186. 10.1016/j.ydbio.2019.12.012.

11. Fink, R.D., and McClay, D.R. (1985). Three cell recognition changes accompany the ingression of sea urchin primary mesenchyme cells. Dev Biol 107, 66–74. 10.1016/0012-1606(85)90376-8.

12. Magie, C.R., Daly, M., and Martindale, M.Q. (2007). Gastrulation in the cnidarian *Nematostella vectensis* occurs via invagination not ingression. Developmental Biology 305, 483–497. 10.1016/j.ydbio.2007.02.044.

13. Kam, Z., Minden, J.S., Agard, D.A., Sedat, J.W., and Leptin, M. (1991). *Drosophila* gastrulation: analysis of cell shape changes in living embryos by three-dimensional fluorescence microscopy. Development 112, 365–370.

14. Leptin, M. (1991). *twist* and *snail* as positive and negative regulators during *Drosophila* mesoderm development. Genes Dev. 5, 1568–1576. 10.1101/gad.5.9.1568.

15. Campos-Ortega, J.A., and Hartenstein, V. (1997). The Embryonic Development of Drosophila melanogaster (Springer) 10.1007/978-3-662-22489-2.

16. Warga, R.M., and Kimmel, C.B. (1990). Cell movements during epiboly and gastrulation in zebrafish. Development 108, 569–580. 10.1242/dev.108.4.569.

17. Godard, B.G., Coolen, M., Le Panse, S., Gombault, A., Ferreiro-Galve, S., Laguerre, L., Lagadec, R., Wincker, P., Poulain, J., Da Silva, C., et al. (2014). Mechanisms of endoderm formation in a cartilaginous fish reveal ancestral and homoplastic traits in jawed vertebrates. Biol Open 3, 1098–1107. 10.1242/bio.20148037.

18. Stower, M.J., Diaz, R.E., Fernandez, L.C., Crother, M.W., Crother, B., Marco, A., Trainor, P.A., Srinivas, S., and Bertocchini, F. (2015). Bi-modal strategy of gastrulation in reptiles. Developmental Dynamics 244, 1144–1157. 10.1002/dvdy.24300.

19. Winklbauer, R. (2020). Mesoderm and endoderm internalization in the *Xenopus* gastrula. Curr Top Dev Biol 136, 243–270. 10.1016/bs.ctdb.2019.09.002.

20. Leptin, M., and Grunewald, B. (1990). Cell shape changes during gastrulation in Drosophila. Development 110, 73–84.

21. Sweeton, D., Parks, S., Costa, M., and Wieschaus, E. (1991). Gastrulation in *Drosophila*: the formation of the ventral furrow and posterior midgut invaginations. Development 112, 775–789.

22. Martin, A.C. (2020). The Physical Mechanisms of *Drosophila* Gastrulation: Mesoderm and Endoderm Invagination. Genetics 214, 543–560. 10.1534/genetics.119.301292.

23. Urbansky, S., González Avalos, P., Wosch, M., and Lemke, S. (2016). Folded gastrulation and T48 drive the evolution of coordinated mesoderm internalization in flies. eLife 5. 10.7554/eLife.18318.

24. Fleig, R., and Sander, K. (1988). Honeybee morphogenesis: embryonic cell movements that shape the larval body. Development 103, 525–534. 10.1242/dev.103.3.525.

25. Benton, M.A., and Pavlopoulos, A. (2014). *Tribolium* embryo morphogenesis. BioArchitecture 4, 16–21. 10.4161/bioa.27815.

26. Pointer, M.D., Gage, M.J.G., and Spurgin, L.G. (2021). *Tribolium* beetles as a model system in evolution and ecology. Heredity 126, 869–883. 10.1038/s41437-021-00420-1.

27. Campbell, J.F., Athanassiou, C.G., Hagstrum, D.W., and Zhu, K.Y. (2022). *Tribolium* castaneum: A Model Insect for Fundamental and Applied Research. Annual Review of Entomology 67, 347–365. 10.1146/annurev-ento-080921-075157.

28. Sommer, R.J., and Tautz, D. (1994). Expression patterns of *twist* and *snail* in *Tribolium* (Coleoptera) suggest a homologous formation of mesoderm in long and short germ band insects. Developmental Genetics 15, 32–37. 10.1002/dvg.1020150105.

29. Handel, K., Basal, A., Fan, X., and Roth, S. (2005). *Tribolium castaneum twist*: gastrulation and mesoderm formation in a short-germ beetle. Dev Genes Evol 215, 13–31. 10.1007/s00427-004-0446-9.

30. Stappert, D., Frey, N., Levetzow, C. von, and Roth, S. (2016). Genome-wide identification of *Tribolium* dorsoventral patterning genes. Development 143, 2443–2454. 10.1242/dev.130641.

31. Benton, M.A., Frey, N., Nunes da Fonseca, R., von Levetzow, C., Stappert, D., Hakeemi, M.S., Conrads, K.H., Pechmann, M., Panfilio, K.A., Lynch, J.A., et al. (2019). Fog signaling has diverse roles in epithelial morphogenesis in insects. eLife 8, e47346. 10.7554/eLife.47346.

32. Pönisch, W., Yanakieva, I., Salbreux, G., and Paluch, E.K. (2024). Cell shape noise strength regulates shape dynamics during EMT-associated cell spreading. Preprint at bioRxiv, 10.1101/2024.10.14.618199 https://doi.org/10.1101/2024.10.14.618199.

33. Patel, N.H. (1994). The evolution of arthropod segmentation: insights from comparisons of gene expression patterns. Dev. Suppl., 201–207.

34. Seher, T.C., and Leptin, M. (2000). Tribbles, a cell-cycle brake that coordinates proliferation and morphogenesis during *Drosophila* gastrulation. Current Biology 10, 623–629. 10.1016/S0960-9822(00)00502-9.

35. Mata, J., Curado, S., Ephrussi, A., and Rørth, P. (2000). Tribbles Coordinates Mitosis and Morphogenesis in *Drosophila* by Regulating String/CDC25 Proteolysis. Cell 101, 511–522. 10.1016/S0092-8674(00)80861-2.

36. Großhans, J., and Wieschaus, E. (2000). A Genetic Link between Morphogenesis and Cell Division during Formation of the Ventral Furrow in *Drosophila*. Cell 101, 523–531. 10.1016/S0092-8674(00)80862-4.

37. Benton, M.A., Akam, M., and Pavlopoulos, A. (2013). Cell and tissue dynamics during *Tribolium* embryogenesis revealed by versatile fluorescence labeling approaches. Development 140, 3210–3220. 10.1242/dev.096271.

38. Dawes-Hoang, R.E., Parmar, K.M., Christiansen, A.E., Phelps, C.B., Brand, A.H., and Wieschaus, E.F. (2005). folded gastrulation, cell shape change and the control of myosin localization. Development 132, 4165–4178. 10.1242/dev.01938.

39. Martin, A.C., Kaschube, M., and Wieschaus, E.F. (2009). Pulsed contractions of an actin–myosin network drive apical constriction. Nature 457, 495–499. 10.1038/nature07522.

40. Martin, A.C., and Goldstein, B. (2014). Apical constriction: themes and variations on a cellular mechanism driving morphogenesis. Development 141, 1987–1998. 10.1242/dev.102228.

41. Simões, S., Oh, Y., Wang, M.F.Z., Fernandez-Gonzalez, R., and Tepass, U. (2017). Myosin II promotes the anisotropic loss of the apical domain during *Drosophila* neuroblast ingression. J Cell Biol 216, 1387–1404. 10.1083/jcb.201608038.

42. Sato, N., Rosa, V.S., Makhlouf, A., Kretzmer, H., Kumar, A.S., Grosswendt, S., Mattei, A.L., Courbot, O., Wolf, S., Boulanger, J., et al. (2024). Basal delamination during mouse gastrulation primes pluripotent cells for differentiation. Developmental Cell 59, 1252–1268.e13. 10.1016/j.devcel.2024.03.008.

43. van Leen, E.V., di Pietro, F., and Bellaïche, Y. (2020). Oriented cell divisions in epithelia: from force generation to force anisotropy by tension, shape and vertices. Current Opinion in Cell Biology 62, 9–16. 10.1016/j.ceb.2019.07.013.

44. Gudipaty, S.A., and Rosenblatt, J. (2017). Epithelial Cell Extrusion: pathways and pathologies. Semin Cell Dev Biol 67, 132–140. 10.1016/j.semcdb.2016.05.010.

45. Box, K., Joyce, B.W., and Devenport, D. (2019). Epithelial geometry regulates spindle orientation and progenitor fate during formation of the mammalian epidermis. eLife 8, e47102. 10.7554/eLife.47102.

46. Lisica, A., Fouchard, J., Kelkar, M., Wyatt, T.P.J., Duque, J., Ndiaye, A.-B., Bonfanti, A., Baum, B., Kabla, A.J., and Charras, G.T. (2022). Tension at intercellular junctions is necessary for accurate orientation of cell division in the epithelium plane. Proceedings of the National Academy of Sciences 119, e2201600119. 10.1073/pnas.2201600119.

47. Breiman, L. (2001). Random Forests. Machine Learning 45, 5–32. 10.1023/A:1010933404324.

48. Sokolova, M., and Lapalme, G. (2009). A systematic analysis of performance measures for classification tasks. Information Processing & Management 45, 427–437. 10.1016/j.ipm.2009.03.002.

49. Lundberg, S., and Lee, S.-I. (2017). A Unified Approach to Interpreting Model Predictions. Preprint at arXiv, 10.48550/arXiv.1705.07874 https://doi.org/10.48550/arXiv.1705.07874.

50. Ponce-Bobadilla, A.V., Schmitt, V., Maier, C.S., Mensing, S., and Stodtmann, S. (2024). Practical guide to SHAP analysis: Explaining supervised machine learning model predictions in drug development. Clin Transl Sci 17, e70056. 10.1111/cts.70056.

51. Edgar, B.A., and O’Farrell, P.H. (1989). Genetic control of cell division patterns in the *Drosophila* embryo. Cell 57, 177–187. 10.1016/0092-8674(89)90183-9.

52. Edgar, B.A., and O’Farrell, P.H. (1990). The three postblastoderm cell cycles of *Drosophila* embryogenesis are regulated in G2 by *string*. Cell 62, 469–480. 10.1016/0092-8674(90)90012-4.

53. Edgar, B.A., Lehman, D.A., and O’Farrell, P.H. (1994). Transcriptional regulation of *string* (*cdc25*): a link between developmental programming and the cell cycle. Development 120, 3131–3143. 10.1242/dev.120.11.3131.

54. Ko, C.S., Kalakuntla, P., and Martin, A.C. (2020). Apical Constriction Reversal upon Mitotic Entry Underlies Different Morphogenetic Outcomes of Cell Division. MBoC 31, 1663–1674. 10.1091/mbc.E19-12-0673.

55. Hertzler, P.L., and Clark, W.H. (1992). Cleavage and gastrulation in the shrimp *Sicyonia ingentis*: invagination is accompanied by oriented cell division. Development 116, 127–140. 10.1242/dev.116.1.127.

56. Gerberding, M. (1997). Germ band formation and early neurogenesis of *Leptodora kindti* (Cladocera): first evidence for neuroblasts in the entomostracan crustaceans. Invertebrate Reproduction & Development 32, 63–73. 10.1080/07924259.1997.9672605.

57. Pawlak, J.B., Sellars, M.J., Wood, A., and Hertzler, P.L. (2010). Cleavage and gastrulation in the Kuruma shrimp *Penaeus* (Marsupenaeus) *japonicus* (Bate): A revised cell lineage and identification of a presumptive germ cell marker. Development, Growth & Differentiation 52, 677–692. 10.1111/j.1440-169X.2010.01205.x.

58. MacBride, E.W. (1898). The Early Development of *Amphioxus*. J Cell Sci s2-40, 589–612. 10.1242/jcs.s2-40.160.589.

59. Mathiah, N., Despin-Guitard, E., Stower, M., Nahaboo, W., Eski, S.E., Singh, S.P., Srinivas, S., and Migeotte, I. (2020). Asymmetry in the frequency and position of mitosis in the mouse embryo epiblast at gastrulation. EMBO reports 21, e50944. 10.15252/embr.202050944.

60. Despin-Guitard, E., Rosa, V.S., Plunder, S., Mathiah, N., Van Schoor, K., Nehme, E., Merino-Aceituno, S., Egea, J., Shahbazi, M.N., Theveneau, E., et al. (2024). Non-apical mitoses contribute to cell delamination during mouse gastrulation. Nat Commun 15, 7364. 10.1038/s41467-024-51638-6.

61. Byrum, C.A. (2001). An analysis of hydrozoan gastrulation by unipolar ingression. Dev Biol 240, 627–640. 10.1006/dbio.2001.0484.

62. Kraus, Y., Flici, H., Hensel, K., Plickert, G., Leitz, T., and Frank, U. (2014). The embryonic development of the cnidarian *Hydractinia echinata*. Evolution & Development 16, 323–338. 10.1111/ede.12100.

63. Kraus, Yu.A., and Markov, A.V. (2017). Gastrulation in Cnidaria: The key to an understanding of phylogeny or the chaos of secondary modifications? Biol Bull Rev 7, 7–25. 10.1134/S2079086417010029.

64. Technau, U. (2020). Gastrulation and germ layer formation in the sea anemone *Nematostella vectensis* and other cnidarians. Mechanisms of Development 163, 103628. 10.1016/j.mod.2020.103628.

65. Vetrova, A.A., Lebedeva, T.S., Saidova, A.A., Kupaeva, D.M., Kraus, Y.A., and Kremnyov, S.V. (2022). From apolar gastrula to polarized larva: Embryonic development of a marine hydroid, *Dynamena pumila*. Developmental Dynamics 251, 795–825. 10.1002/dvdy.439.

66. Mansour, K. (1927). The Development of the larval and adult mid-gut of *Calandra oryzae* (Linn.): The Rice Weevil. J Cell Sci s2*-*71, 313–352. 10.1242/jcs.s2-71.282.313.

67. Davis, C. (1967). A comparative study of larval embryogenesis in the mosquito *Culex fatigans* Wiedemann (Diptera : Culicidae) and the sheep-fly *Lucilia sericata* Meigen (Diptera : Calliphoridae). Aust J Zool 15, 547–579. 10.1071/ZO9670547.

68. Anderson, D.T. (1969). On the embryology of the cirripede crustaceans *Tetraclita rosea* (Krauss), *Tetraclita purpurascens* (Wood), *Chthamalus antennatus* (Darwin) and *Chamaesipho columna* (Spengler) and some considerations of crustacean phylogenetic relationships. Philosophical Transactions of the Royal Society of London. B, Biological Sciences 256, 183–235. 10.1098/rstb.1969.0041.

69. Alwes, F., Hinchen, B., and Extavour, C.G. (2011). Patterns of cell lineage, movement, and migration from germ layer specification to gastrulation in the amphipod crustacean *Parhyale hawaiensis*. Developmental Biology 359, 110–123. 10.1016/j.ydbio.2011.07.029.

70. Murakami, M.S., Moody, S.A., Daar, I.O., and Morrison, D.K. (2004). Morphogenesis during *Xenopus* gastrulation requires Wee1-mediated inhibition of cell proliferation. Development 131, 571–580. 10.1242/dev.00971.

71. Davis, G.K., and Patel, N.H. (2002). Short, long, and beyond: molecular and embryological approaches to insect segmentation. Annu Rev Entomol 47, 669–699. 10.1146/annurev.ento.47.091201.145251.

72. Clark, E. (2017). Dynamic patterning by the *Drosophila* pair-rule network reconciles long-germ and short-germ segmentation. PLOS Biology 15, e2002439. 10.1371/journal.pbio.2002439.

73. Gadagkar, S.R., and Newfeld, S.J. (2026). Selection for rapid embryonic development sculpted the new gene Shrew into an accelerator of *Drosophila* dorsal-ventral axis specification. Genetics 233, iyag087. 10.1093/genetics/iyag087.

74. Vellutini, B.C., Cuenca, M.B., Krishna, A., Szałapak, A., Modes, C.D., and Tomancak, P. (2025). Patterned invagination prevents mechanical instability during gastrulation. Nature 646, 627–636. 10.1038/s41586-025-09480-3.

75. Dey, B., Kaul, V., Kale, G., Scorcelletti, M., Takeda, M., Wang, Y.-C., and Lemke, S. (2025). Divergent evolutionary strategies pre-empt tissue collision in gastrulation. Nature 646, 637–646. 10.1038/s41586-025-09447-4.

76. Tozluoǧlu, M., and Mao, Y. (2020). On folding morphogenesis, a mechanical problem. Philosophical Transactions of the Royal Society B: Biological Sciences 375, 20190564. 10.1098/rstb.2019.0564.

77. Johnston, L.A. (2000). Cell cycle: The trouble with *tribbles*. Current Biology 10, R502–R504. 10.1016/S0960-9822(00)00559-5.

78. Nelsen, O.E. (1934). The segregation of the germ cells in the grasshopper, *Melanoplus differentialis* (acrididae; orthoptera). Journal of Morphology 55, 545–575. 10.1002/jmor.1050550306.

79. Thomas, A.J. (1936). The Embryonic Development of the Stick-Insect, Carausius morosus. J Cell Sci s2*-*78, 487–511. 10.1242/jcs.s2-78.311.487.

80. Roonwal, M.L. (1937). Studies on the Embryology of the African Migratory Locust, *Locusta migratoria migratorioides* Reiche and Frm. (Orthoptera, Acrididae). II. Organogeny. Philosophical Transactions of the Royal Society of London. Series B, Biological Sciences 227, 175–244.

81. Ullmann, S.L. (1964). The origin and structure of the mesoderm and the formation of the coelomic sacs in *Tenebrio molitor* L. [Insecta, Coleoptera]. Philosophical Transactions of the Royal Society of London. Series B, Biological Sciences 248, 245–277. 10.1098/rstb.1964.0012.

82. Ruberson, J.R., Larsen, J.R., and Jorgensen, C.D. (1987). Embryogenesis of the Codling Moth, *Cydia pomonella* (Lepidoptera: Tortricidae). Ann Entomol Soc Am 80, 561–570. 10.1093/aesa/80.5.561.

83. Bucher, G. (2009). Tribolium castaneum beetle book (Online release).

84. Sullivan, W., Ashburner, M., and Hawley, R.S. (2000). Drosophila Protocols pp. 150–151, Chapter 9 (CSHL Press).

85. Choi, H.M.T., Schwarzkopf, M., Fornace, M.E., Acharya, A., Artavanis, G., Stegmaier, J., Cunha, A., and Pierce, N.A. (2018). Third-generation in situ hybridization chain reaction: multiplexed, quantitative, sensitive, versatile, robust. Development 145, dev165753. 10.1242/dev.165753.

86. Clark, E., Battistara, M., and Benton, M.A. (2022). A timer gene network is spatially regulated by the terminal system in the *Drosophila* embryo. eLife 11, e78902. 10.7554/eLife.78902.

87. Tidswell, O.R.A., Benton, M.A., and Akam, M. (2021). The neuroblast timer gene nubbin exhibits functional redundancy with gap genes to regulate segment identity in *Tribolium*. Development 148, dev199719. 10.1242/dev.199719.

88. Shaner, N.C., Lambert, G.G., Chammas, A., Ni, Y., Cranfill, P.J., Baird, M.A., Sell, B.R., Allen, J.R., Day, R.N., Israelsson, M., et al. (2013). A bright monomeric green fluorescent protein derived from *Branchiostoma lanceolatum*. Nat Methods 10, 407–409. 10.1038/nmeth.2413.

89. Bindels, D.S., Haarbosch, L., van Weeren, L., Postma, M., Wiese, K.E., Mastop, M., Aumonier, S., Gotthard, G., Royant, A., Hink, M.A., et al. (2017). mScarlet: a bright monomeric red fluorescent protein for cellular imaging. Nat. Methods 14, 53–56. 10.1038/nmeth.4074.

90. Pfeiffer, B.D., Ngo, T.-T.B., Hibbard, K.L., Murphy, C., Jenett, A., Truman, J.W., and Rubin, G.M. (2010). Refinement of Tools for Targeted Gene Expression in *Drosophila*. Genetics 186, 735–755. 10.1534/genetics.110.119917.

91. Suzuki, T., Ito, M., Ezure, T., Kobayashi, S., Shikata, M., Tanimizu, K., and Nishimura, O. (2006). Performance of expression vector, pTD1, in insect cell-free translation system. Journal of Bioscience and Bioengineering 102, 69–71. 10.1263/jbb.102.69.

92. Pfeiffer, B.D., Truman, J.W., and Rubin, G.M. (2012). Using translational enhancers to increase transgene expression in *Drosophila*. PNAS 109, 6626–6631. 10.1073/pnas.1204520109.

93. van Oers, M.M., Vlak, J.M., Voorma, H.O., and Thomas, A.A.M. (1999). Role of the 3’ untranslated region of baculovirus p10 mRNA in high-level expression of foreign genes. J Gen Virol 80 ( *Pt 8*), 2253–2262. 10.1099/0022-1317-80-8-2253.

94. Jain, A., Ulman, V., Mukherjee, A., Prakash, M., Cuenca, M.B., Pimpale, L.G., Münster, S., Haase, R., Panfilio, K.A., Jug, F., et al. (2020). Regionalized tissue fluidization is required for epithelial gap closure during insect gastrulation. Nat Commun 11, 5604. 10.1038/s41467-020-19356-x.

95. Brown, J.E., Klement, J.F., and McAllister, W.T. (1986). Sequences of three promoters for the bacteriophage SP6 RNA polymerase. Nucleic Acids Res 14, 3521–3526. 10.1093/nar/14.8.3521.

96. Benton, M.A. (2018). A revised understanding of *Tribolium* morphogenesis further reconciles short and long germ development. PLoS Biol 16, e2005093. 10.1371/journal.pbio.2005093.

97. Schindelin, J., Arganda-Carreras, I., Frise, E., Kaynig, V., Longair, M., Pietzsch, T., Preibisch, S., Rueden, C., Saalfeld, S., Schmid, B., et al. (2012). Fiji: an open-source platform for biological-image analysis. Nature Methods 9, 676–682. 10.1038/nmeth.2019.

98. Stringer, C., Wang, T., Michaelos, M., and Pachitariu, M. (2021). Cellpose: a generalist algorithm for cellular segmentation. Nat Methods 18, 100–106. 10.1038/s41592-020-01018-x.

99. Van Rossum, G., and Drake, F.L. (2009). Python 3 Reference Manual (Python Software Foundation).

100. Legland, D., Arganda-Carreras, I., and Andrey, P. (2016). MorphoLibJ: integrated library and plugins for mathematical morphology with ImageJ. Bioinformatics 32, 3532–3534. 10.1093/bioinformatics/btw413.

101. Inc, T.M. (2022). MATLAB version: 9.13.0 (R2022b). (The MathWorks Inc.).

102. Chung, M.K., and Taylor, J. (2004). Diffusion smoothing on brain surface via finite element method. In 2004 2nd IEEE International Symposium on Biomedical Imaging: Nano to Macro (IEEE Cat No. 04EX821), pp. 432–435 Vol. 1. 10.1109/ISBI.2004.1398567.

103. Rousseeuw, P.J., Ruts, I., and Tukey, J.W. (1999). The Bagplot: A Bivariate Boxplot. The American Statistician 53, 382–387. 10.1080/00031305.1999.10474494.

104. Verboven, S., and Hubert, M. (2005). LIBRA: a MATLAB library for robust analysis. Chemometrics and Intelligent Laboratory Systems 75, 127–136. 10.1016/j.chemolab.2004.06.003.

105. Fuentes, M.A., and He, B. (2022). The cell polarity determinant Dlg1 facilitates epithelial invagination by promoting tissue-scale mechanical coordination. Development 149, dev200468. 10.1242/dev.200468.

106. Sommer, C., Straehle, C., Köthe, U., and Hamprecht, F.A. (2011). ilastik: Interactive Learning and Segmentation.

107. Berg, S., Kutra, D., Kroeger, T., Straehle, C.N., Kausler, B.X., Haubold, C., Schiegg, M., Ales, J., Beier, T., Rudy, M., et al. (2019). ilastik: interactive machine learning for (bio)image analysis. Nat Methods 16, 1226–1232. 10.1038/s41592-019-0582-9.

108. Harris, C.R., Millman, K.J., van der Walt, S.J., and others (2020). Array programming with NumPy. Nature 585, 357–362. 10.1038/s41586-020-2649-2.

109. McKinney, W. (2010). Data structures for statistical computing in Python. In Proceedings of the 9th Python in Science Conference, pp. 51–56. 10.25080/Majora-92bf1922-00a.

110. Virtanen, P., Gommers, R., Oliphant, T.E., Haberland, M., Reddy, T., Cournapeau, D., Burovski, E., Peterson, P., Weckesser, W., Bright, J., et al. (2020). SciPy 1.0: fundamental algorithms for scientific computing in Python. Nat Methods 17, 261–272. 10.1038/s41592-019-0686-2.

111. van der Walt, S., Schönberger, J.L., Nunez-Iglesias, J., Boulogne, F., Warner, J.D., Yager, N., Gouillart, E., and Yu, T. (2014). scikit-image: image processing in Python. PeerJ 2, e453. 10.7717/peerj.453.

112. Hunter, J.D. (2007). Matplotlib: A 2D Graphics Environment. Computing in Science & Engineering 9, 90–95. 10.1109/MCSE.2007.55.

113. Waskom, M.L. (2021). Seaborn: statistical data visualization. Journal of Open Source Software 6, 3021. 10.21105/joss.03021.

114. Herbert, S., Valon, L., Mancini, L., Dray, N., Caldarelli, P., Gros, J., Esposito, E., Shorte, S.L., Bally-Cuif, L., Aulner, N., et al. (2021). LocalZProjector and DeProj: a toolbox for local 2D projection and accurate morphometrics of large 3D microscopy images. BMC Biol 19, 136. 10.1186/s12915-021-01037-w.

115. Aigouy, B., and Prud’homme, B. (2022). Segmentation and Quantitative Analysis of Epithelial Tissues. Methods Mol Biol 2540, 387–399. 10.1007/978-1-0716-2541-5_20.

116. Lundberg, S.M., Erion, G., Chen, H., DeGrave, A., Prutkin, J.M., Nair, B., Katz, R., Himmelfarb, J., Bansal, N., and Lee, S.-I. (2020). From local explanations to global understanding with explainable AI for trees. Nat Mach Intell 2, 56–67. 10.1038/s42256-019-0138-9.

